# The heterogeneous distribution of glial cell marker proteins in the myelinated region of the normal rat optic nerve

**DOI:** 10.1101/2025.03.19.643607

**Authors:** June Kawano

**Affiliations:** Department of Gene Therapy and Regenerative Medicine, Kagoshima University Graduate School of Medical and Dental Sciences, Kagoshima, 890-8544, Japan; Division of Morphological Sciences, Department of Neurology, Kagoshima University Graduate School of Medical and Dental Sciences, Kagoshima, 890-8544, Japan; Laboratory for Neuroanatomy, Department of Neurology, Kagoshima University Graduate School of Medical and Dental Sciences, Kagoshima, 890-8544, Japan

**Author notes:** Correspondence to: June Kawano, M.D., Ph.D., Department of Gene Therapy and Regenerative Medicine, Kagoshima University Graduate School of Medical and Dental Sciences, 35-1, Sakuragaoka 8-chome, Kagoshima, 890-8544, Japan.

**Keywords:** glial fibrillary acidic protein, glutamine synthetase, myelin basic protein, myelin debris-like particles, immunohistochemistry, RRID: AB_2109645, RRID: AB_259853, RRID: AB_397879, RRID: AB_10120129, RRID: AB_509997, RRID: AB_94266

## Abstract

Glial cells play a critical role in the maintenance of neuronal activity in the optic nerve. The present study reports the distribution of glial structural proteins (GFAP: glial fibrillary acidic protein; MBP: myelin basic protein), a glial functional protein (GS: glutamine synthetase), and of a cell nuclear marker (bisBenzimide) in the various myelinated regions of the normal rat optic nerve. Fourteen optic nerves from 12 male Sprague-Dawley rats were used. Immunohistochemistry and confocal microscopy were employed to investigate the distribution of GFAP, MBP, GS, and of bisBenzimide along the longitudinal plane of the myelinated region. GFAP-immunoreactivity and GS-immunoreactivity were strong in the distal (anterior)-most part but weak in the proximal (posterior) part, demonstrating a significant decrease in strength along the longitudinal plane of the myelinated region. bisBenzimide labeling was also strong in the distal-most part but weak in the proximal part, indicating a significant difference in strength across the myelinated region. MBP-immunoreactive particles and cell nuclei were densely distributed in the distal-most part; however, they were sparsely dispersed in the proximal part, showing a significant difference. The heterogeneous distribution of GFAP, GS, bisBenzimide, cell nuclei, and of MBP-immunoreactive particles along the longitudinal plane may represent an important functional adaptation reflecting the non-uniform nature of the physiological and structural environment of the myelinated region. Notably, the concentrations of GFAP, GS, and of MBP-immunoreactive particles in the distal-most part of the myelinated region suggest that this area is under physiologically stressed conditions in the normal rat optic nerve.

**Key points:**

- The present study reports the distribution of glial structural proteins (GFAP: glial fibrillary acidic protein; MBP: myelin basic protein), and of a glial functional protein (GS: glutamine synthetase) in the various myelinated regions of the normal rat optic nerve.
- GFAP-immunoreactivity and GS-immunoreactivity were strong in the distal (anterior)-most part but weak in the proximal (posterior) part.
- MBP-immunoreactive particles were densely distributed in the distal-most part; however, they were sparsely dispersed in the proximal part.
- These results suggest that the distal-most part is not under physiological but rather under physiologically stressed conditions in the normal rat optic nerve.

**Graphical Abstract:**

The present study reports the distribution of glial structural pro-teins (GFAP: glial fibrillary acidic protein; MBP: myelin basic protein), and of a glial functional protein (GS: glutamine syn-thetase) in the various myelinated regions of the normal rat optic nerve.
GF AP-immunoreactivity and GS-immunoreactivity were strong in the distal (anterior)-most part but weak in the proximal (poste-rior) part.
MBP-immunoreactive particles were densely distributed in the distal-most part; however, they were sparsely dispersed in the proximal part.
These results suggest that the distal-most part is not under physi-ological but rather under physiologically stressed conditions in the normal rat optic nerve.

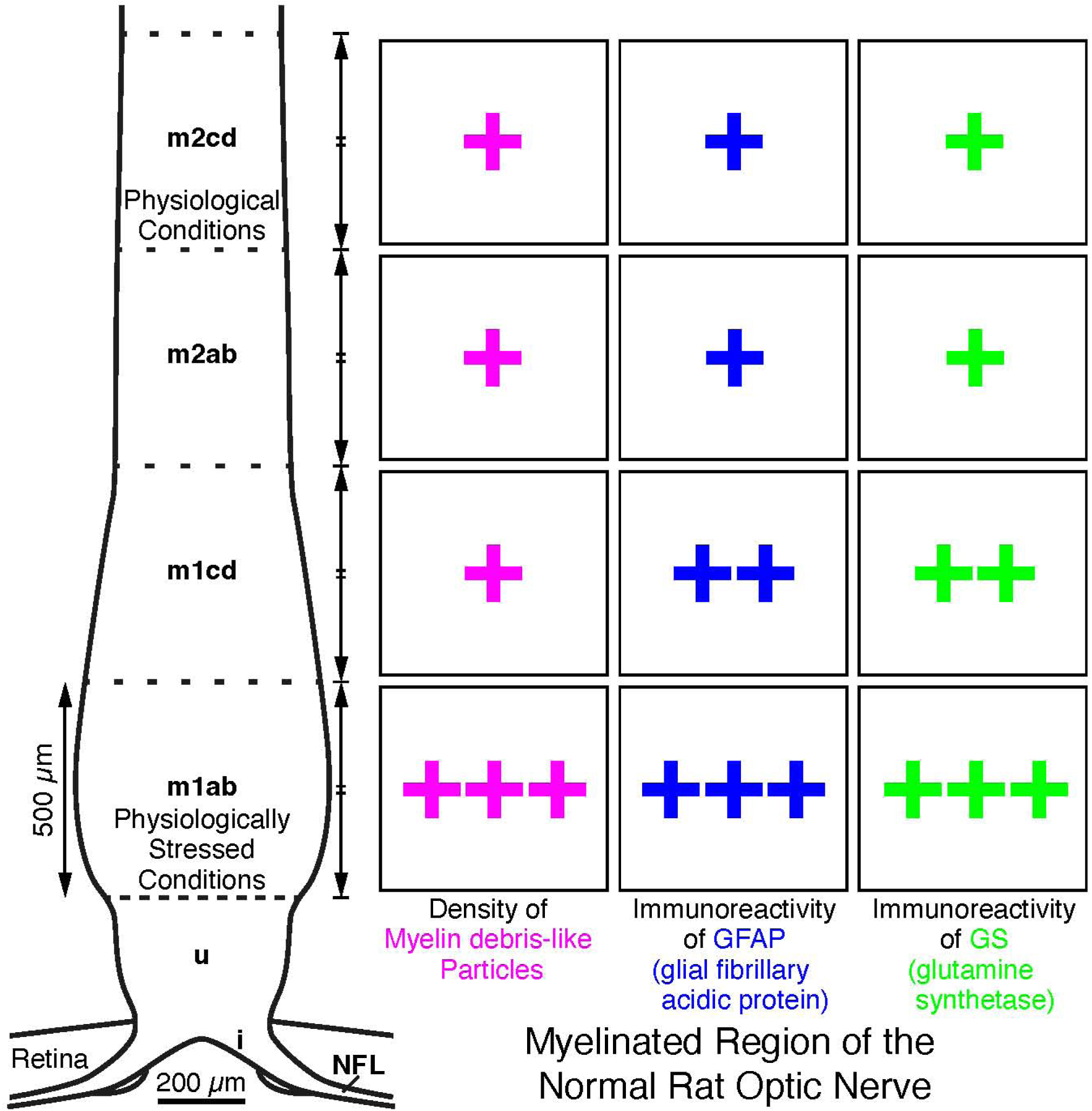

## 1 Introduction

Glial cells play a critical role in the maintenance of neuronal activity in the optic nerve. The optic nerve comprises the axons of retinal ganglion cells, interstitial glial cells (astrocytes, oligodendrocytes, neuron glial (NG2) cells, and microglia), and the fibrovascular septa of the pia mater (Yazdankhah et al. 2021). These glial cells are essential components of the optic nerve, collaborating to sustain its neuronal activity and structural stability. In the myelinated region, oligodendrocytes form myelin sheaths around axons, facilitating rapid, metabolically efficient conduction of action potentials, and are essential for axon integrity and nodal function (Yazdankhah et al. 2021). Astrocytes were originally considered to be passive structural elements, but it is now evident that they are dynamic and multifunctional cells that are important for potassium homeostasis, metabolic processes, and axon-glial signaling, as well as the physiology of nodes of Ranvier, and for contributions to the blood-brain barrier (Khakh and Deneen 2019; Yazdankhah et al. 2021). In addition to interacting with axons, glial cells constantly interact with each other, thereby forming a robust and highly efficient network that supports the functionality of the optic nerve. Importantly, since the optic nerve does not contain any synapses but consists solely of axons that extend the entire length of the nerve, it has become a popular model for studying the interactions of glial cells with axons without the need to consider their complex roles in maintaining synapse functionality (Yazdankhah et al. 2021).

The optic nerve is essentially a tract in the brain, the fiber of which has no neurolemma, and is enclosed by meninges, which is different from any peripheral nerve (Bron et al. 1997). The axons of retinal ganglion cells are unmyelinated in the retina of the rat and become myelinated when they leave the eye (Black et al. 1985; Kawano 2015). The chemoarchitecture of the rodent optic nerve has been investigated by several research groups, focusing on the distributions of astrocytic filaments and myelinated axons (Ffrench-Constant et al. 1988; Morcos and Chan-Ling 2000; Howell et al. 2007; Kawano et al. 2008; Sun et al. 2009; Balaratnasingam et al. 2014; Kawano 2015). A very high density of astrocytic filaments has been observed at the junction of the retina and optic nerve. The posterior limit of these astrocytic filaments primarily coincides with the beginning of axon myelination (Morcos and Chan-Ling 2000; Kawano 2015). Consequently, the rodent optic nerve has been divided into at least three regions: the intraretinal (prelaminar), unmyelinated (laminar, lamina cribrosa, or astrocytic filament dense), and myelinated (retrolaminar, retrobulbar, or astrocytic filament sparse) regions (May 2003; Morrison et al. 2005; Howell et al. 2007; Kawano et al. 2008; Balaratnasingam et al. 2014; Kawano 2015).

Until recently, investigations of the chemoarchitecture of the rodent optic nerve have been progressed by using immunohistochemistry. In the myelinated region of the normal rodent optic nerve, bundles of myelinated axons have been shown to interdigitate with astrocytes (Morcos and Chan-Ling 2000; Kawano 2015). A quantitative study demonstrated that the chemoarchitecture of glial fibrillary acidic protein (GFAP) in the myelinated region is heterogeneously organized: GFAP immunoreactivity is strong in the distal (anterior) region; however, it is moderate in the proximal (posterior) region (Kawano 2015). Regarding glutamine synthetase (GS), a quantitative study demonstrated that the chemoarchitecture of GS in the myelinated region is also heterogeneously organized: GS immunoreactivity is moderate in the distal region, while it is weak in the proximal region. The majority of GS-immunoreactive glial cells in the myelinated region are oligodendrocytes (Kawano 2015). On the myelinated axon, the nodes of Ranvier are distributed in a regularly spaced manner. Nearly all (>95%) of the nodes contain processes from astrocytes, while fewer than half of the nodes are contacted by both the NG2 cell and astrocyte processes in the adult rat optic nerve. Ultrastructural analysis revealed that NG2 cells extend fine, finger-like projections that often contact both the nodal axolemma and the outermost paranodal myelin loops, while astrocytes extend broader processes covering the entire nodal gap, suggesting different roles of the two glial cell types at the node (Serwanski et al. 2017). In the companion article (Kawano 2026), we demonstrated the distribution of myelin basic protein (MBPh^1^)-immunoreactive particles in the distal-most part of the myelinated region of the normal rat optic nerve. We confirmed that MBPh-immunoreaction to these particles was authentic. Additionally, we showed that the morphological characteristics of MBPh-immunoreactive particles were similar to those of myelin debris, suggesting that the site of the distribution was under physiologically stressed conditions in the normal rat optic nerve (Weinger et al. 2011; Zhou et al. 2019; Wang et al. 2020; Yao et al. 2023; Kawano 2026). Thus, the necessary neuroanatomical facts, demonstrated by using immunohistochemistry in the myelinated region of the rodent optic nerve, have been accumulated. However, information on the detailed distribution patterns of glial cell proteins and of glial cell DNA remains limited, especially regarding the detailed distribution pattern of MBPh-immunoreactive particles. Furthermore, the shape descriptors of MBPh-immunoreactive particles in each myelinated region remain unknown.

Here we show, by using quantitative immunohistochemical analyses, the concentrations of GFAP, GS, cell nuclei, and of MBPh-immunoreactive particles in the distal-most part of the myelinated region. Additionally, we demonstrate that the shape descriptors of MBPh-immunoreactive particles remain primarily stable from the distal myelinated region through the proximal myelinated region. Finally, based on these observations, we propose that this site of concentrations is not in physiological conditions but rather under physiologically stressed conditions in the normal rat optic nerve, since myelin debris-like MBPh-immunoreactive particles accumulate at this site (Weinger et al. 2011; Zhou et al. 2019; Wang et al. 2020; Yao et al. 2023; Kawano 2026). Furthermore, the assertion that this site is under physiologically stressed conditions is reinforced by two results obtained as outlined below. The concentration of GFAP at this site suggests reactive astrogliosis, a secondary event following insults in the central nervous system (CNS; Messam et al. 2002; McAteer and Choudhury 2009; Sofroniew 2009). Given that GS in oligodendrocytes increases in chronic pathological conditions in both mice and humans (Ben Haim et al. 2021), a significant increase in GS immunoreactivity among GS-immunoreactive cells at this site indicates that this site is under physiologically stressed conditions.

## 2 Materials and Methods

### 2.1 Animals and tissue preparation

Male rats (n=12; 12 weeks old; Slc:SD; CLEA Japan, Tokyo, Japan) were deeply anesthetized with sodium pentobarbital (50 mg/kg, i.p.) and perfused transcardially with 4 % paraformaldehyde dissolved in 0.1 M sodium phosphate buffer (PB; pH 7.4) at 4°C. The eyeballs, including the optic nerve, were removed from the skull, stored in the same fixative for 48 hours, and then immersed in 30 % saccharose in 0.1 M PB at 4°C until they sank. The eyeballs, including the optic nerve, were frozen in powdered dry ice and sectioned in the meridian plane at a thickness of 25 µm on a cryostat. Sections were collected in a cryoprotectant medium (Warr et al. 1981; 33.3% saccharose, 1% polyvinylpyrrolidone (K-30), and 33.3% ethylene glycol in 0.067M sodium phosphate buffer (pH 7.4) containing 0.067% sodium azide) and stored at –30 °C prior to use (Kawano et al. 2008; Kawano 2015).

All animal experiments were approved by the Institutional Animal Care and Use Committee of Kagoshima University (MD11112; MD15029; MD18058), and were conducted according to the related guidelines and applicable laws in Japan.

### 2.2 Antibody characterization

Please refer to Table 1 for a list of all antibodies used. These antibodies are cataloged in the “Journal of Comparative Neurology antibody database (Version 14)” except for the rabbit anti-glutamine synthetase (GS) antibody.

**TABLE 1.**
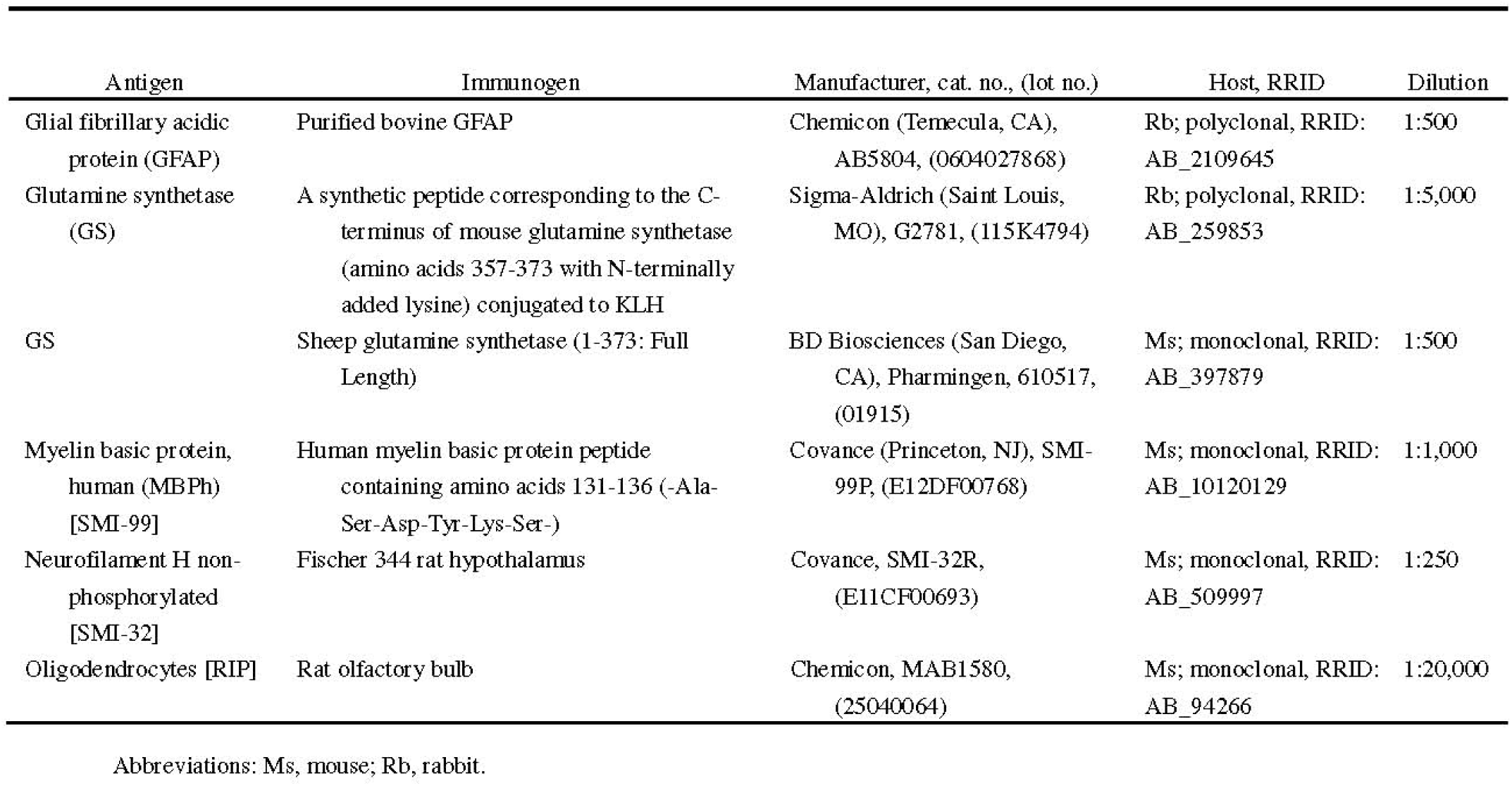
Primary antibodies used in this study.

#### Glial fibrillary acidic protein (GFAP)

Anti-GFAP rabbit serum (Chemicon, Temecula, CA, USA) stains a single protein band of 51 kDa in the mouse brain on immunoblots (Jalil et al. 2005; Kawano, unpublished observations). This serum is also specific for GFAP on immunoblots in extracts from the mouse spinal cord (Darman et al. 2004) and rat retina (Chang et al. 2007). Astrocytes are immunolabeled with this antibody against GFAP (Morcos and Chan-Ling 2000). The staining obtained with this anti-GFAP antibody in the rat was similar to published results on the rat (Saari et al. 1997; Morcos and Chan-Ling 2000; Ju et al. 2005; Chang et al. 2007; Kawano 2015).

#### Glutamine synthetase (GS)

The rabbit anti-GS antibody (Sigma-Aldrich, Saint Louis, MO, USA) recognizes a single protein band of 45 kDa in extracts from the rat brain. Immunoblotting analysis demonstrates that the staining of GS is specifically inhibited by the GS immunizing peptide (amino acids 357-373 with N-terminally added lysine). This amino acid sequence is identical in human, bovine, rat, hamster, and pig GS, and is highly conserved in chicken GS (single amino acid substitution; manufacturer’s technical information).

The mouse monoclonal anti-GS antibody (clone 6; BD Biosciences Pharmingen, San Diego, CA, USA) visualizes a single protein band (45 kDa) in extracts from the rat brain on immunoblots (manufacturer’s technical information). Müller cells in the retina and glial cells in the optic nerve are labeled with these antibodies against GS (Riepe and Norenburg 1977).

The staining obtained with the two antibodies in the rat was similar to that previously reported in the mouse (Haverkamp and Wässle 2000; Hojo et al. 2000; Kawano et al. 2008) and the rat (Riepe and Norenburg 1977; Zabouri et al. 2011; Kawano 2015).

#### Myelin basic protein, human (MBPh)

The mouse monoclonal antibody against the human myelin basic protein (clone SMI-99; Covance, Princeton, NJ, USA) detects four bands between 14 and 21 kDa, corresponding to four myelin basic protein (MBP) isoforms on immunoblots of the mouse cerebellum (Dyer et al. 1996; Talos et al. 2006). The SMI-99 antibody detects MBP from most mammalian species. The pig and chicken MBP do not react, while the guinea pig MBP has slight reactivity. This antibody does not react with the 14kDa form of the rat MBP. The SMI-99 antibody detects the developing and adult myelin, and distinguishes oligodendrocytes from astrocytes, microglia, neurons and other cells in brain sections (manufacturer’s technical information). The staining obtained with this antibody against the human myelin basic protein was similar to that previously reported in the rat (Morcos and Chan-Ling 2000), except for the concentration of MBPh-immunoreactive particles (Kawano 2026).

#### Neurofilament H non-phosphorylated

The mouse monoclonal antibody against neurofilament H non-phosphorylated (clone SMI-32; Covance, Princeton, NJ, USA) visualizes two bands (200 and 180 kDa) on conventional immunoblots, merging into a single neurofilament H line on two-dimensional blots (Sternberger et al. 1982; Goldstein et al. 1983; Goldstein et al. 1987). The SMI-32 antibody reacts with a non-phosphorylated epitope from neurofilament H found in most mammalian species. The reaction is masked when the epitope is phosphorylated (Sternberger and Sternberger 1983). The SMI-32 antibody immunocytochemically visualizes neuronal cell bodies, dendrites, and some thick axons in the central and peripheral nervous systems, whereas thin axons are not revealed (manufacturer’s technical information). The staining obtained with this SMI-32 antibody was similar to published results on the mouse (Howell et al. 2007) and the rat (Kawano 2015).

#### Oligodendrocytes

Mature oligodendrocytes are immunostained with the mouse monoclonal antibody against oligodendrocytes [RIP] (clone NS-1; Chemicon; Friedman et al. 1989). The specificity of this antibody is strictly verified in the rat CNS (central nervous system; Friedman et al. 1989). Strong RIP staining is primarily confined to the white matter in the rat cerebellum, while moderate RIP staining is distributed in the granular layer. RIP staining is hardly detected in the molecular layer, which is composed of unmyelinated parallel fibers (Friedman et al. 1989). The staining obtained with this antibody against oligodendrocytes [RIP] was similar to that previously reported in the rat (Saari et al. 1997; Morcos and Chan-Ling 2000; Kawano 2015).

### 2.3 Immunohistochemistry

Sections were processed using double-label immunohistochemistry as previously described (Kawano et al. 2008; Kawano 2015). Free-floating sections were pre-incubated for 2 hours with a 10% normal goat serum (NGS) blocking solution at 4 °C and then immunoreacted for 4 days with a mixture of rabbit and mouse primary antibodies in a 10% NGS blocking solution at 4 °C (Table 1). After two rinses for 10 minutes in 0.02M phosphate buffered saline (PBS) containing 0.3% Triton X-100 (PBST), the sections were incubated with a mixture of two secondary antibodies in PBS containing 5% NGS and 0.3% Triton X-100 for 24 hours at 4 °C. The two secondary antibodies used were Alexa Fluor 488 conjugated with the F(ab’)_2_ fragment of goat anti-rabbit IgG (H+L) (1:200; Molecular Probes, Eugene, OR, USA) and Alexa Fluor 594 conjugated to the F(ab’)_2_ fragment of goat anti-mouse IgG (H+L) (1:200; Molecular Probes). The sections were washed once for 10 minutes in PBST, and then twice in PBS. The sections were mounted onto hydrophilic silanized slides (Dako Japan, Tokyo, Japan) by using an equal-parts mixture of a 0.6% gelatin solution and PBS. After being air-dried, the sections were subjected to nuclear staining by using a bisBenzimide (bBM; Hoechst 33258, Sigma-Aldrich; 0.1 mg/ml) solution and coverslipped with VECTASHIELD mounting medium (Vector Laboratories, Burlingame, CA, USA).

To eliminate the possibility of any cross-reaction between the secondary and primary antibodies from the different species, one of the two primary antibodies was removed. No cross-reactivity was observed in these control experiments (data not shown).

In all cases, each staining protocol was conducted on a minimum of 5 optic nerves from 5 separate rats. GFAP staining was executed on a total of 11 optic nerves from 10 separate rats. GS staining was performed on 6 optic nerves from 6 separate rats. MBPh staining was conducted on 5 optic nerves from 5 separate rats. bBM staining was executed on a total of 14 optic nerves from 12 separate rats.

### 2.4 Photomicrographs

Fluorescent photomicrographs were obtained using an LSM700 confocal laser scanning microscope (Carl Zeiss Jena GmbH, Jena, Germany) at the Joint Research Laboratory, Kagoshima University Graduate School of Medical and Dental Sciences. The images were subsequently transferred to Adobe Photoshop CS5 (Adobe Systems, San Jose, CA, USA) for processing. The brightness and contrast of the images were adjusted. No other adjustments were made.

### 2.5 Image analysis

The quantitation of all images was performed using ImageJ2 (Version 2.9.0/1.53t; developed by Wayne Rasband, National Institute of Mental Health, Bethesda, MD, USA).

### 2.6 Quantitative morphological analyses in each myelinate region

#### 2.6.1 Measurement of the area of each region, and that of the mean pixel intensity of GFAP immunoreactivity in each myelinated region

Confocal microscopic images of GFAP immunoreactivity were assembled using a photomerge function in Adobe Photoshop CS5 (Layout; Reposition; Figure 1A, B). Measurements were conducted in the shaded area of each myelinated region depicted in Figure 1C and 1D. The area is also shown in GFAP immunoreactivity image of each myelinated region in Figure 3. Before taking measurements, the color channels were split to isolate the green channel, and the background was removed from the measurement area (Figure 3). For each image, the process executed by the ImageJ2 program included the following: (a) setting the threshold at 1-255 to binarize the image (Manual); (b) analyzing particles to measure area and mean pixel intensity (size, 100 µm^2^-infinity; circularity, 0.00-1.00; check “Clear results” and “Add to manager”).

**FIGURE 1.**
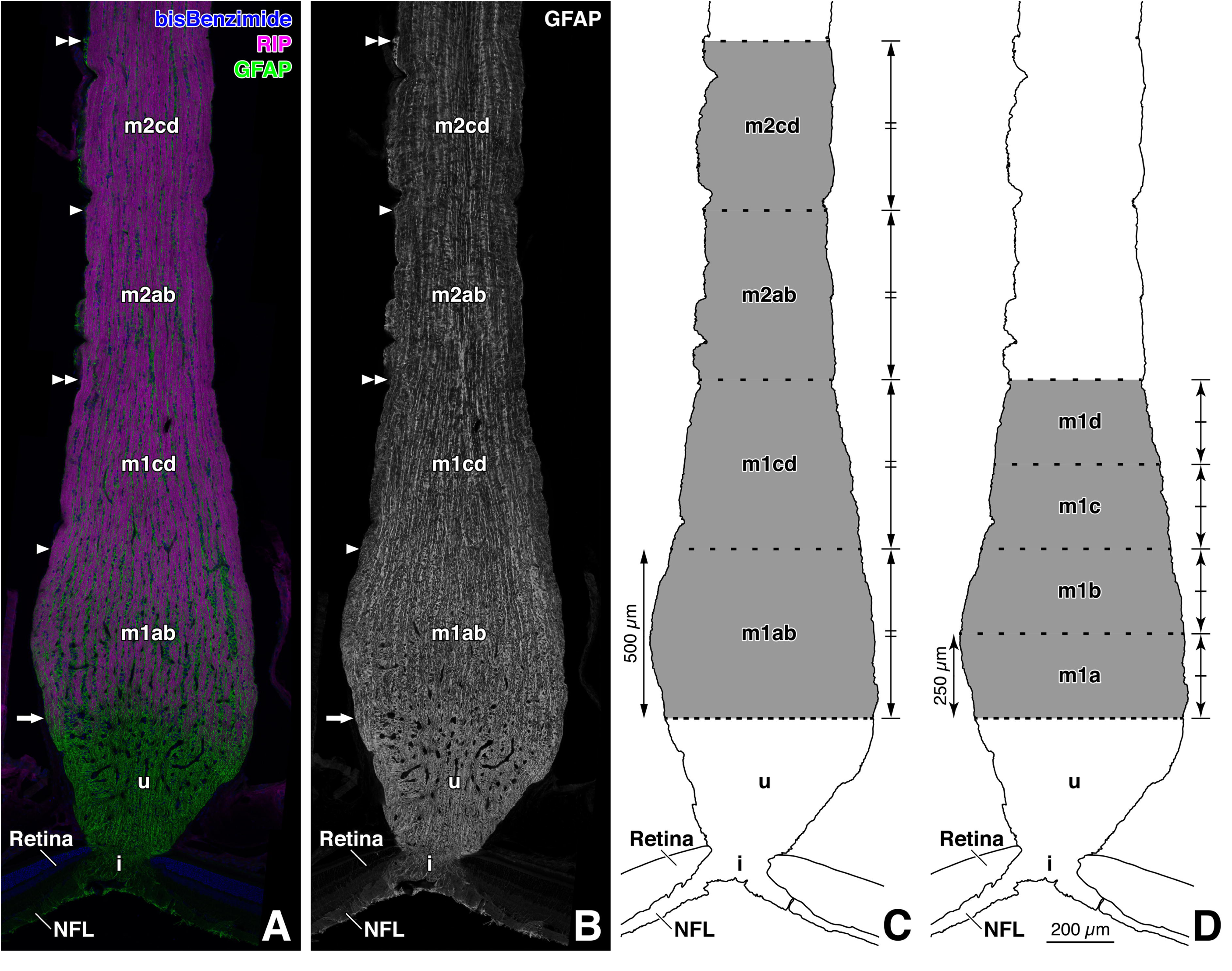
**A-B:** Distribution of astrocytes and oligodendrocytes in the normal rat optic nerve. The panels show a longitudinal section through the paramedian part and represent double immunostaining for an astrocyte marker (glial fibrillary acidic protein, GFAP; **B**; Alexa Fluor 488 label; green in **A**) and for an oligodendrocyte marker (RIP; Alexa Fluor 594 label; magenta in **A**). Cell nuclei were labeled with bisBenzimide (Hoechst 33258; blue in **A**). Panel **A** is an overlaid image. The arrows indicate the border between the unmyelinated (u) and myelinated regions. The double arrowheads represent boundaries between the subregions: the boundary between myelinated regions 1cd (m1cd) and 2ab (m2ab), or the boundary between myelinated regions 2cd (m2cd) and 3. The arrowheads show borders between the sub-subregions: the border between myelinated region 1ab (m1ab) and m1cd, or the border between m2ab and m2cd. Panel **A** is a photo montage composed of 10 square fluorescent images. These images were taken with an LSM 700 confocal microscope (Carl Zeiss, Jena, Germany). Certain parts of the background were filled with black color. Note that GFAP immunoreactivity in the m1ab region was stronger than that in the remaining three myelinated regions: the m1cd, m2ab, and m2cd regions. Interestingly, columns of astrocytes were observed to interdigitate with myelinated (RIP-immunoreactive) nerve fiber bundles in the m1ab region. i, intraretinal region; NFL, nerve fiber layer of the retina. **C:** The diagram showing a division scheme in the rat optic nerve used in this study. In this diagram, shading indicates areas where quantitative analyses were conducted. The optic nerve was divided into six regions: intraretinal (i) region, unmyelinated (u) region, and myelinated regions 1ab (m1ab), 1cd (m1cd), 2ab (m2ab), and 2cd (m2cd). Dashed lines composed of dashes and shorter gaps delineate the border between the u and m1ab regions. In contrast, dashed lines with longer gaps demarcate the boundaries between each subregion: for example, the boundary between the m1ab and m1cd regions, or that between the m1cd and m2ab regions. The one up-down arrow positioned to the left of the m1ab region indicates that the longitudinal length of this region is 500 µm. Additionally, the four up-down arrows positioned to the right of the m1ab, m1cd, m2ab, and m2cd regions, which are of equal length, also show that each region is 500 µm long. Note that the m1ab region was thicker than the remaining three myelinated regions: the m1cd, m2ab, and m2cd regions. NFL, nerve fiber layer of the retina. **D:** The diagram showing another division scheme in the rat optic nerve used in this study. In this diagram, shading shows areas where quantitative analyses were executed. The optic nerve was divided into six regions: intraretinal (i) region, unmyelinated (u) region, and myelinated regions 1a (m1a), 1b (m1b), 1c (m1c), and 1d (m1d). Dashed lines composed of dashes and shorter gaps delineate the border between the u and m1a regions. In contrast, the dashed lines with longer gaps demarcate the boundaries between subregions: for example, the boundary between the m1a and m1b regions, or that between the m1d region and myelinated region 2. The one up-down arrow positioned to the left of the m1a region indicates that the longitudinal length of this is 250 µm. The four up-down arrows positioned to the right of the m1a, m1b, m1c, and m1d regions, which are of equal length, also show that each region is 250 µm long. NFL, nerve fiber layer of the retina. Scale bar = 200 µm at the bottom of **D** for **A-C**.

#### 2.6.2 Measurement of the area and the mean pixel intensity of GFAP-immunoreactive filaments in each myelinated region

For each image, the process executed by the ImageJ2 program included the following: (a) establishing the threshold in order to binarize the image (Auto (Default; IJ_IsoData) or Manual); (b) analyzing particles to measure the area and mean pixel intensity of GFAP immunoreactive filaments (size, 0 µm^2^-infinity; circularity, 0.00-1.00; check “Clear Results” and “Add to manager”).

The mean pixel intensity of GFAP-immunoreactive filaments in each myelinated region was determined as a weighted average, manually calculated based on the outputs of the aforementioned measurement processes (the mean pixel intensity of each GFAP-immunoreactive filament and the area of each GFAP-immunoreactive filament).

#### 2.6.3 Measurement of the proportion of the total area of GFAP-immunoreactive filaments in each myelinated region

The proportion of the total area of GFAP-immunoreactive filaments was manually calculated based on the total of area of each GFAP-immunoreactive filament and on the area of each myelinated region.

#### 2.6.4 Measurement of the area of each myelinated region, and that of the mean pixel intensities of GS immunoreactivity, bBM labeling, and of MBPh immunoreactivity in each myelinated region

Confocal microscopic images of GS immunoreactivity, bBM labeling, and of MBPh immunoreactivity were manually assembled using Adobe Photoshop CS5 (Figure 2). Measurements were conducted in the shaded area of each myelinated region depicted in Figure 1C and 1D. The areas where the measurements were taken are also shown in the GS immunoreactivity images of each myelinated region in Figure 4, in the bBM labeling images of each myelinated region in Figure 5, and in the MBPh immunoreactivity images of each myelinated region in Figure 6. Before the measurements, the color channels were split to obtain the green channel for the GS immunoreactivity image, the blue channel for the bBM labeling image, and the red channel for the MBPh immunoreactivity image; subsequently, the background was removed from the measurement area (Figures 4–6). For each image, the process executed by the ImageJ2 program included the following: (a) setting the threshold at 1-255 in order to binarize the image (Manual); (b) analyzing particles to measure area and mean pixel intensity (size, 1.0 µm^2^-infinity; circularity, 0.00-1.00; check “Clear results” and “Add to manager”).

**FIGURE 2.**
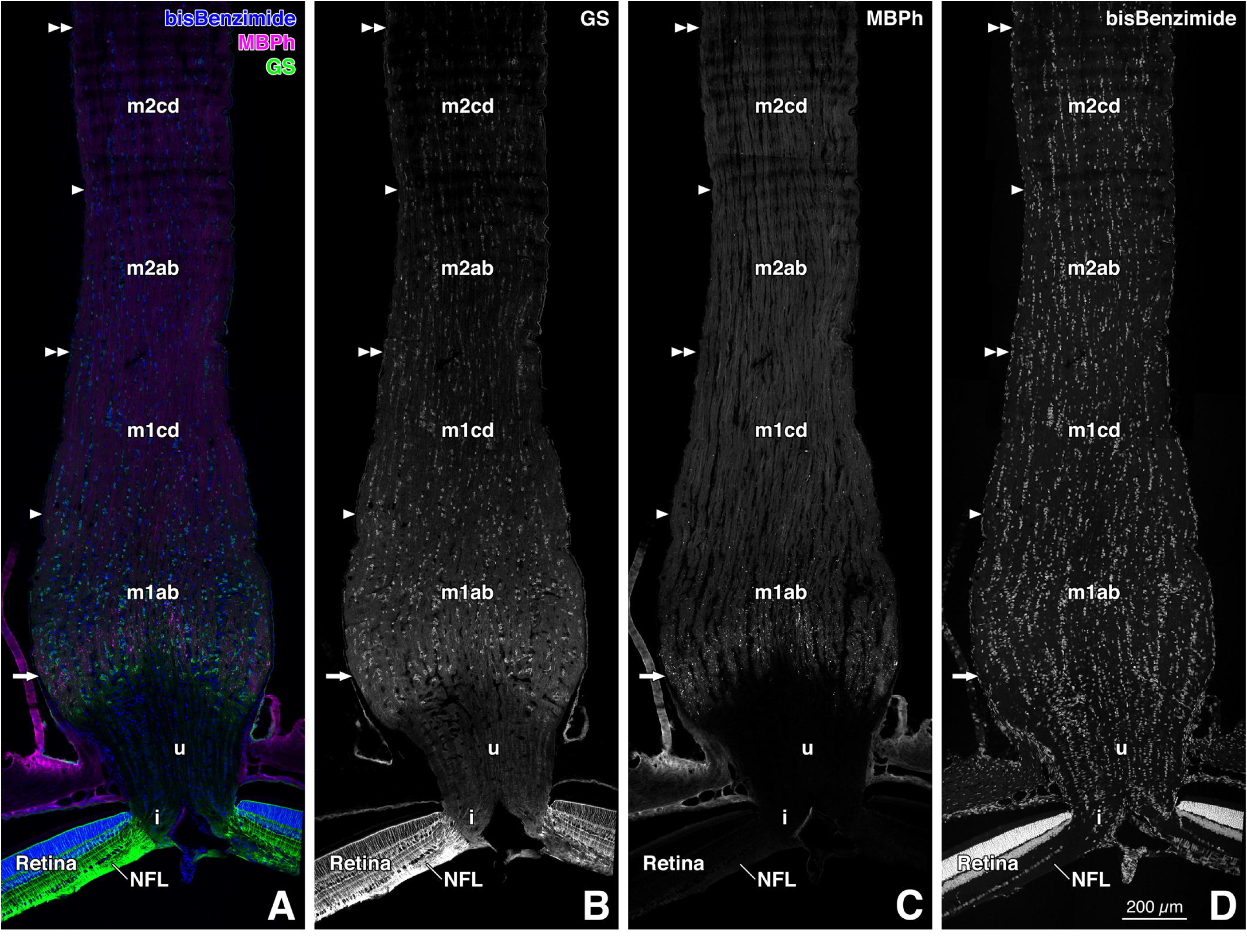
Distribution of glutamine synthetase (GS)-immunoreactive glial cells, myelin basic protein (MBPh^3^)-immunoreactive nerve fiber bundles, and of MBPh-immunoreactive particles in the normal rat optic nerve. The panels show a longitudinal section through the paramedian part and represent double immunostaining for a glial cell marker (glutamine synthetase, GS; **B**; Alexa Fluor 488 label; green in **A**) and for a myelin marker (myelin basic protein, MBPh; **C**; Alexa Fluor 594 label; magenta in **A**). Cell nuclei were labeled with bisBenzimide (Hoechst 33258; **D**; blue in **A**). Panel **A** is an overlaid image. The arrows indicate the border between the unmyelinated (u) region and myelinated region 1ab (m1ab). The double arrowheads represent boundaries between the subregions: the boundary between myelinated regions 1cd (m1cd) and 2ab (m2ab), or the boundary between myelinated regions 2cd (m2cd) and 3. The arrowheads show borders between the sub-subregions: the border between the m1ab and m1cd regions, or the border between the m2ab and m2cd regions. Panel **A** is a photo montage composed of 11 square fluorescent images. These images were taken with an LSM 700 confocal microscope (Carl Zeiss, Jena, Germany). Certain parts of the background in panel **A** were filled with black color. Note that many MBPh-immunoreactive particles were distributed in the m1ab region, especially in the distal (anterior) half. Interestingly, the distal half of the m1ab region was the major site for GS distribution in the normal rat optic nerve; however, GS immunoreactivity in this region was markedly weaker than that in the retina. NFL, nerve fiber layer of the retina. Scale bar = 200 µm at the bottom of **D** for **A-C**.

**FIGURE 3.**
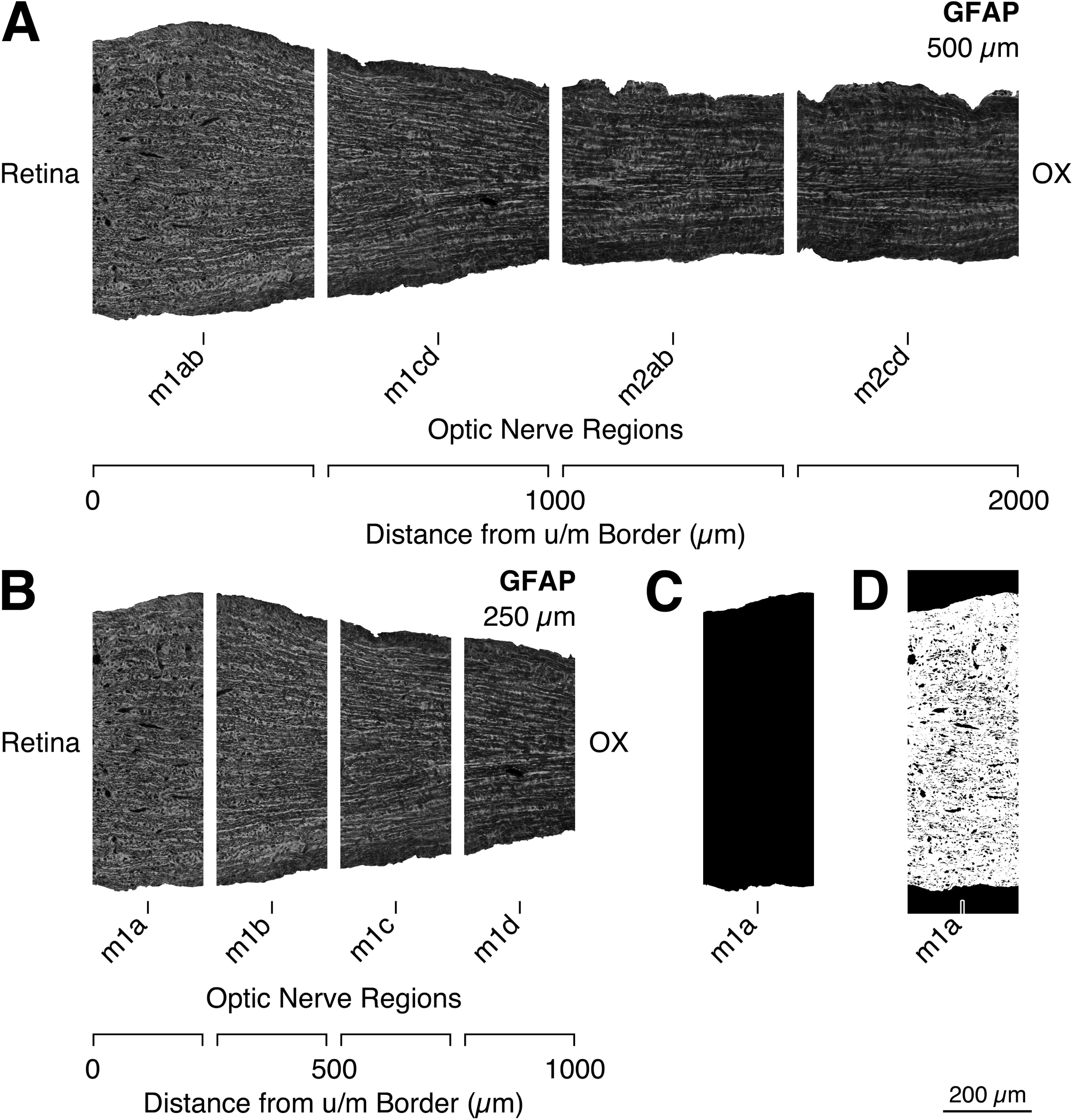
**A-B:** Images used for quantitative analyses of glial fibrillary acidic protein (GFAP) distribution along the course of the myelinated regions of the normal rat optic nerve. The original image of Figure 1B, in which brightness and contrast were not adjusted (meaning the brightness levels of the image histogram were not moved or stretched), was modified to create these images. The black background of the original image was manually removed using Adobe Photoshop CS5. The left edges of the m1ab (myelinated region 1ab) and m1a (myelinated region 1a) images indicate the border between the unmyelinated and myelinated regions. The horizontal length of each image is 500 µm in **A** and 250 µm in **B**. The background color of each modified image is completely white for presentation purposes.^4^ The “Retina” and the “OX (optic chiasm)” show the orientation of each modified image. The scale at the bottom of each modified image represents a distance from the border between the unmyelinated and myelinated regions (Distance from u/m Border). **C:** The black binary image used to measure the area of the m1a region. **D:** The white parts of this binary image indicate ROIs (regions of interest) used to analyze mean pixel intensity of GFAP immunoreactivity and the proportion of the total area occupied by GFAP-immunoreactive filaments in the m1a region. m1b, myelinated region 1b; m1c, myelinated region 1c; m1cd, myelinated region 1cd; m1d, myelinated region 1d; m2ab, myelinated region 2ab; m2cd, myelinated region 2cd. Scale bar = 200 µm in the lower right corner of **D** for **A-C**.

**FIGURE 4.**
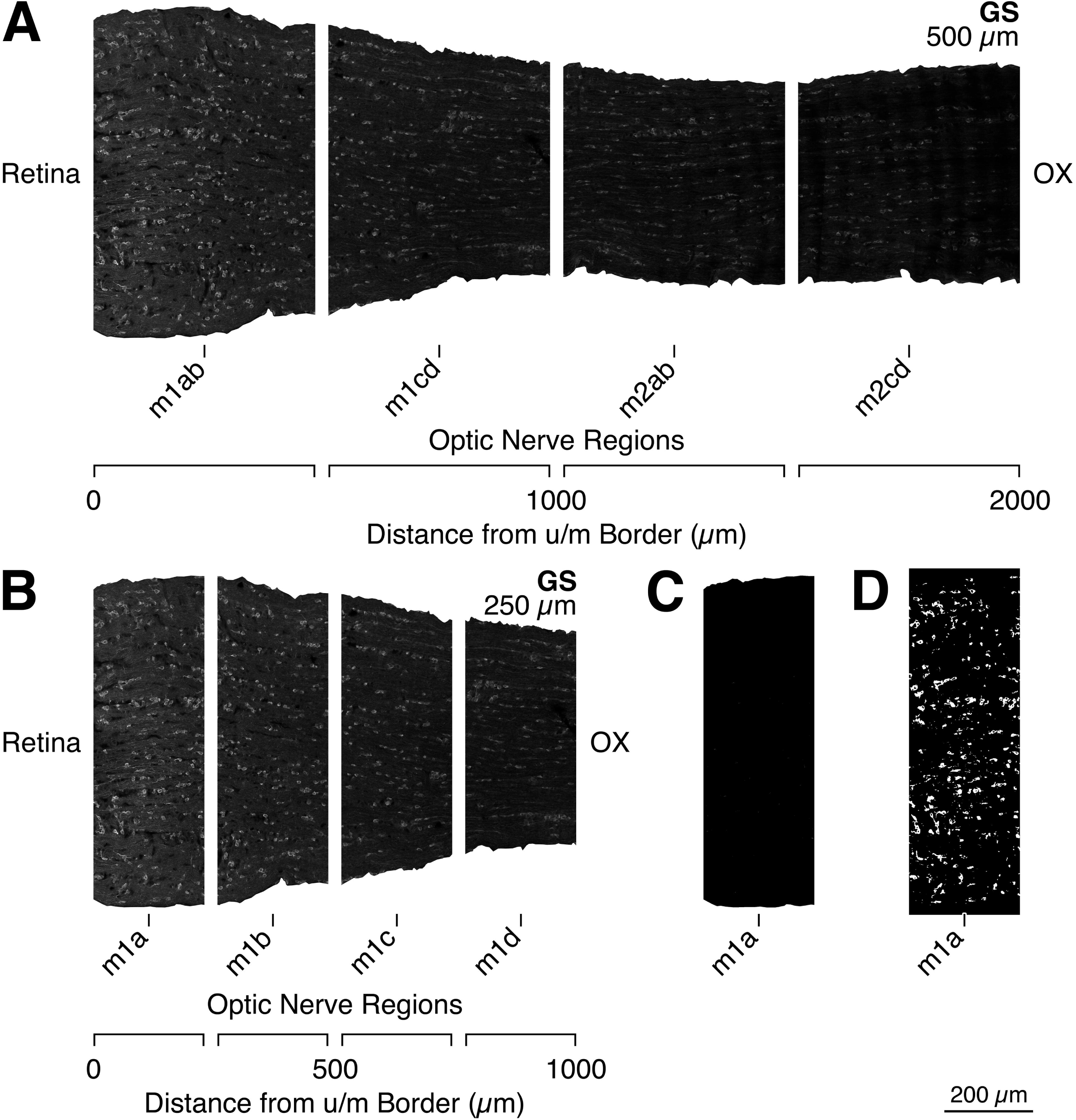
**A-B:** Images used for quantitative analyses of glutamine synthetase (GS) distribution along the course of the myelinated regions of the normal rat optic nerve. The original image of Figure 2B, in which brightness and contrast were not adjusted (meaning the brightness levels of the image histogram were not moved or stretched), was modified to create these images. The black background of the original image was manually removed using Adobe Photoshop CS5. The left edges of the m1ab (myelinated region 1ab) and m1a (myelinated region 1a) images indicate the border between the unmyelinated and myelinated regions. The horizontal length of each image is 500 µm in **A** and 250 µm in **B**. The background color of each modified image is completely white for presentation purposes (see footnote 4 in the legend for Figure 3). The “Retina” and the “OX (optic chiasm)” show the orientation of each modified image. The scale at the bottom of each modified image represents the distance from the border between the unmyelinated and myelinated regions (Distance from u/m Border). **C:** The black binary image used to measure the area of the m1a region. **D:** The white parts of this binary image indicate ROIs (regions of interest) used to analyze the mean pixel intensity of GS immunoreactivity, the proportion of total area, density, and the mean area of GS-immunoreactive cells in the m1a region. m1b, myelinated region 1b; m1c, myelinated region 1c; m1cd, myelinated region 1cd; m1d, myelinated region 1d; m2ab, myelinated region 2ab; m2cd, myelinated region 2cd. Scale bar = 200 µm in the lower right of **D** for **A-C**.

**FIGURE 5.**
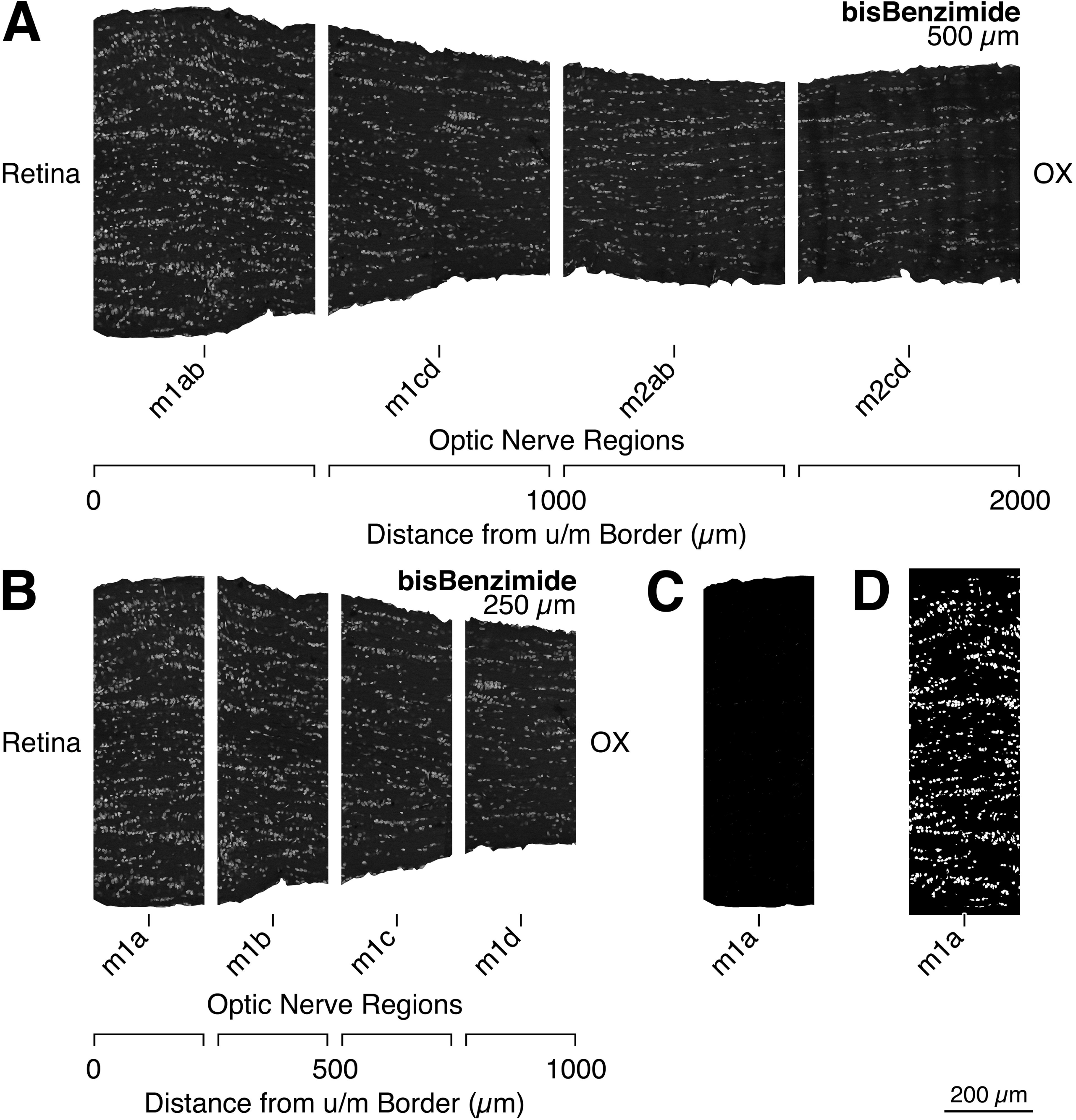
**A-B:** Images used for quantitative analyses of bisBenzimide (bBM) distribution along the course of the myelinated regions of the normal rat optic nerve. The original image of Figure 2D, in which brightness and contrast were not adjusted (meaning the brightness levels of the image histogram were not moved or stretched), was modified to create these images. The black background of the original image was manually removed using Adobe Photoshop CS5. The left edges of the m1ab (myelinated region 1ab) and m1a (myelinated region 1a) images indicate the border between the unmyelinated and myelinated regions. The horizontal length of each image is 500 µm in **A** and 250 µm in **B**. The background color of each modified image is completely white for presentation purposes (see footnote 4 in the legend for Figure 3). The “Retina” and the “OX (optic chiasm)” show the orientation of each modified image. The scale at the bottom of each modified image represents the distance from the border between the unmyelinated and myelinated regions (Distance from u/m Border). **C:** The black binary image used to measure the area of the m1a region. **D:** The white parts of this binary image indicate ROIs (regions of interest) used to analyze the mean pixel intensity of bBM labeling, the proportion of total area, density, and the mean area of bBM-labeled cell nuclei in the m1a region. m1b, myelinated region 1b; m1c, myelinated region 1c; m1cd, myelinated region 1cd; m1d, myelinated region 1d; m2ab, myelinated region 2ab; m2cd, myelinated region 2cd. Scale bar = 200 µm in the lower right corner of **D** for **A-C**.

**FIGURE 6.**
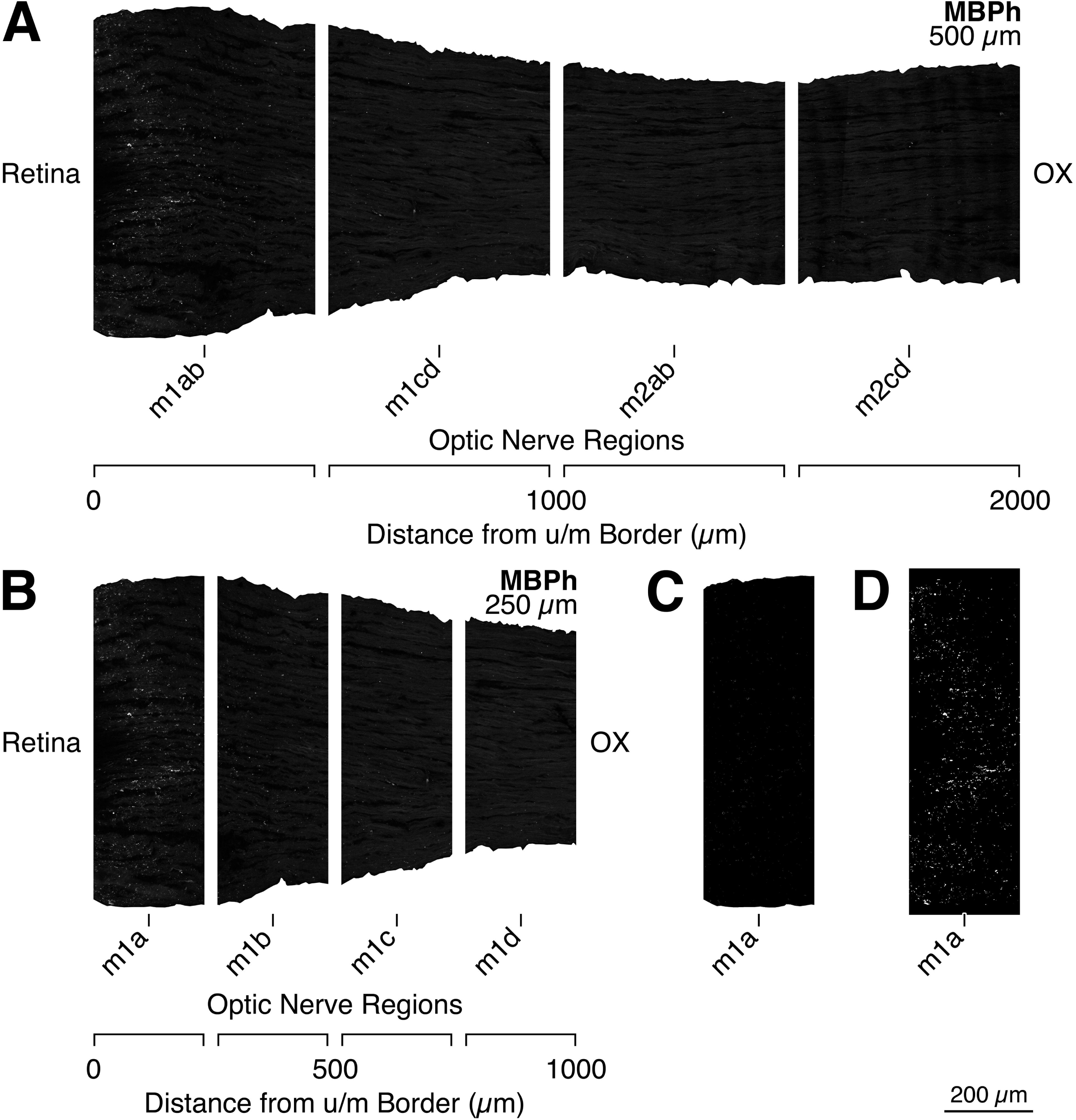
**A-B:** Images used for quantitative analyses of myelin basic protein (MBPh^5^) distribution and for those of MBPh-immunoreactive particles along the course of the myelinated regions of the normal rat optic nerve. The original image of Figure 2C, in which brightness and contrast were not adjusted (meaning the brightness levels of the image histogram were not moved or stretched), was modified to create these images. The black background of the original image was manually removed using Adobe Photoshop CS5. The left edges of the m1ab (myelinated region 1ab) and m1a (myelinated region 1a) images indicate the border between the unmyelinated and myelinated regions. The horizontal length of each image is 500 µm in A and 250 µm in B. The background color of each modified image is completely white for presentation purposes (see footnote 4 in the legend for Figure 3). The “Retina” and the “OX (optic chiasm)” show the orientation of each modified image. The scale at the bottom of each modified image represents the distance from the border between the unmyelinated and myelinated regions (Distance from u/m Border). **C:** The black binary image used to measure the area of the m1a region. **D:** The white parts of this binary image indicate ROIs (regions of interest) used to analyze the mean pixel intensity of MBPh immunoreactivity, the proportion of total area, density, mean area, mean perimeter, and shape descriptors (mean circularity, mean AR (aspect ratio), mean roundness, and mean solidity) of MBPh-immunoreactive particles in the m1a region. m1b, myelinated region 1b; m1c, myelinated region 1c; m1cd, myelinated region 1cd; m1d, myelinated region 1d; m2ab, myelinated region 2ab; m2cd, myelinated region 2cd. Scale bar = 200 µm in the lower right corner of **D** for **A-C**.

#### 2.6.5 Counts of GS-immunoreactive cells

GS-immunoreactive cells were counted in the shaded area of each region depicted in Figure 1C and 1D. The brightness and contrast of images were adjusted for cell count by using Photoshop CS5. A Photoshop image file was opened in Adobe Illustrator CS5 (Adobe Systems) by using the Place function. A small closed circle was inscribed above each soma of the GS immunoreactive cells on the upper layer in Illustrator CS5. These small closed circles were subsequently counted to obtain the number of GS-immunoreactive cells by selecting the circles; the number of the circles was displayed in the Objects item of the Document Info panel in Illustrator CS5. The morphological characteristics of GS-immunoreactive cells were as follows: 1) GS immunoreactivity was clearly seen in the cell body of the GS-immunoreactive cell; 2) The area of the cell nucleus was encircled by strong-to-moderate GS immunoreactivity; 3) Weak GS immunoreactivity was seen in the area of the cell nucleus; 4) All parts of the cell body were located in the measurement area.

#### 2.6.6 Measurement of the mean area and the mean pixel intensity of GS-immunoreactive cells

For each image, the process executed by the ImageJ2 program included the following: (a) conducting the Subtract Background function to remove smooth continuous background from the image (Rolling Ball Algorithm; Radius: 50.0 pixels); (b) performing the Median filter function to reduce noise in the image (Radius: 2.0 pixels); (c) executing the Mean filter function to smooth the image (Radius: 2.0 pixels); (d) establishing the threshold to binarize the image (Auto (Default; IJ_IsoData) or Manual); (e) analyzing particles to measure the area and mean pixel intensity of GS-immunoreactive cells (size: 1.0 µm^2^-infinity; circularity: 0.00-1.00; check “Clear Results”, “Add to manager”, “Exclude on edges” and “Include holes”).

The mean area of GS-immunoreactive cells in each myelinated region was manually calculated based on the outputs of the aforementioned measurement processes (the total area of GS-immunoreactive cells in each myelinated region) and on the counting processes (the total number of GS-immunoreactive cells in each myelinated region).

The mean pixel intensity of GS-immunoreactive cells in each region was determined as a weighted average, manually calculated based on the outputs from the aforementioned measurement processes (the area and the mean pixel intensity of each GS-immunoreactive cell).

#### 2.6.7 Measurement of the mean area, mean pixel intensity, and number of bBM-labeled cell nuclei

For each image, the process executed by the ImageJ2 program included the following: (a) conducting the Subtract Background function to remove smooth continuous background from the image (Rolling Ball Algorithm; Radius: 50.0 pixels); (b) performing the Median filter function to reduce noise in the image (Radius: 2.0 pixels); (c) executing the Mean filter function to smooth the image (Radius: 2.0 pixels); (d) establishing the threshold in order to binarize the image (Auto; Default; IJ_IsoData); (e) applying the watershed; (f) analyzing particles to measure the area and mean pixel intensity of each bBM-labeled cell nucleus and to count bBM-labeled cell nuclei (size: 7.0 µm^2^-infinity; circularity: 0.00-1.00; check “Clear Results”, “Add to manager”, “Exclude on edges” and “Include holes”).

The mean area of bBM-labeled cell nuclei in each myelinated region was manually calculated based on the outputs of the aforementioned measurement processes (the area of each bBM-labeled cell nucleus and the number of bBM-labeled cell nuclei).

The mean pixel intensity of bBM-labeled cell nuclei in each myelinated region was determined as a weighted average, manually calculated based on the outputs of the aforementioned measurement processes (the area and mean pixel intensity of each bBM-labeled cell nucleus).

#### 2.6.8 Measurement of the mean pixel intensity, mean area, mean perimeter, mean shape descriptors, and number of MBPh-immunoreactive particles

For each image, the process executed by the ImageJ2 program included the following: (a) establishing the threshold in order to binarize the image (Manual); (b) analyzing particles to obtain mean pixel intensity, area, perimeter, and shape descriptors (Circularity; AR (aspect ratio); Roundness; Solidity) of each MBPh-immunoreactive particle, as well as counting number of MBPh-immunoreactive particles (size: 1.0 µm^2^-infinity; circularity 0.00-1.00; check “Clear results”, “Add to manager”, and “Exclude on edges”).

The mean pixel intensity of MBPh-immunoreactive particles in each myelinated region was determined as a weighted average, manually calculated based on the outputs of the aforementioned measurement processes (the area and mean pixel intensity of each MBPh-immunoreactive particle).

The mean area, mean perimeter, and mean shape descriptors of MBPh-immunoreactive particles were manually calculated based on the outputs of the aforementioned measurement processes (the area, perimeter, Circularity, AR, Roundness, and Solidity of each MBPh-immunoreactive particle, and the number of MBPh-immunoreactive particles).

#### 2.6.9 Measurement of the proportion of total areas of GS-immunoreactive cells, bBM-labeled cell nuclei, and of MBPh-immunoreactive particles

The proportion of the total areas of GS-immunoreactive cells, bBM-labeled cell nuclei, and of MBPh-immunoreactive particles was manually calculated based on the output from the measurement processes of the area of each GS-immunoreactive cell, bBM-labeled cell nucleus, and of each MBPh-immunoreactive particle on the area of each myelinated region.

#### 2.6.10 Measurement of the density of GS-immunoreactive cells, bBM-labeled cell nuclei, and of MBPh-immunoreactive particles

The density of GS-immunoreactive cells, bBM-labeled cell nuclei, and of MBPh-immunoreactive particles was manually calculated based on the outputs of the counting processes and on the area of each myelinated region.

### 2.7 Statistical analysis

No statistical methods were used to predetermine group sample size. However, our group sample sizes were similar to those in previously published studies by our group and others (Melo et al. 2006; Balaratnasingam et al. 2009; Kawano 2015). All biological replicates (n) are derived from at least five independent experiments. Unless otherwise specified, no data were excluded from analysis. Individual experiment values are represented in box-and-whisker plots. In these box-and-whisker plots, the thick horizontal line in the box indicates the median. The upper and lower borders of the box represent the interquartile range. The upper whisker shows the maximum value, while the lower whisker indicates the minimum value of the distribution.

All statistical analyses were conducted by using EZR (Easy R; Version 1.55; Saitama Medical Center, Jichi Medical University, Saitama, Japan), which serves as a graphical user interface for R (Version 4.1.3; The R Foundation for Statistical Computing, Vienna, Austria). More precisely, it is a modified version of R Commander (Version 2.7-2; Fox 2017) that is designed to add statistical functions frequently used in biostatistics (Kanda 2013). Data were analysed using the Kruskal-Wallis test followed by the Steel-Dwass test for multiple comparisons. A p-value of < 0.05 was considered statistically significant.

## 3 Results

### 3.1 The division of the normal rat optic nerve

A definition of the optic nerve division was essential for the description and interpretation of experimental results; therefore, we outlined our criteria for determining the border of each region of the optic nerve. The criteria principally followed our previously reported rat criteria, except for the subregions in the myelinated region (Kawano 2015).

As shown in Figure 1A, the rat optic nerve in the orbit was divided into three regions: the intraretinal (i), unmyelinated (u), and myelinated (m) regions. This classification was defined by the position of the sensory retina, the distribution of oligodendrocytes (myelinated nerve fibers), and by the strength of GFAP immunoreactivity. The border between the i and u regions was established at the boundary between the sensory retina and the retinal pigment epithelium. Very strong GFAP immunoreactivity was observed in the u region. The border between the u and m regions was defined by the distal limit of myelinated nerve fibers (Figure 1A). This border principally coincided with the boundary defined by the proximal limit of very strong GFAP immunoreactivity; however, protrusions of myelinated nerve fibers were observed in the posterior part of the u region (Figure 1A, B). The border between the nerve fiber layer of the retina (NFL) and the optic nerve proper was established at the medial margin of the area exhibiting strong GS immunoreactivity in the NFL (Kawano 2015; Figure 2A, B).

### 3.2 Division and general overview of the myelinated (m) region in the normal rat optic nerve

The m region was further divided into two subregions: myelinated regions 1 (m1) and 2 (m2; Figure 1). The longitudinal length of each subregion was established at 1mm. Consequently, the boundary between the m1 and m2 regions was determined at the line where the distance from the border between the unmyelinated and myelinated regions (distance from the u/m border) measured 1mm. The proximal boundary of the m2 region was set at the line where the distance from the u/m border measured 2 mm. The m1 region was divided into four sub-subregions: myelinated regions 1a (m1a), 1b (m1b), 1c (m1c), and 1d (m1d). The longitudinal length of each of these sub-subregions was 250 µm (Figure 1D). The m1 region was also divided into two sub-subregions: myelinated regions 1ab (m1ab) and 1cd (m1cd). The longitudinal length of each of these sub-subregions was 500 µm (Figure 1C). The m2 region was also divided into two sub-subregions: myelinated regions 2ab (m2ab) and 2cd (m2cd). Each sub-subregion in the m2 region had a longitudinal length of 500 µm (Figure 1C).

The m1ab region was thicker than the other three myelinated regions: the m1cd, m2ab, and m2cd regions (Figures 1–2).

Densely packed myelinated (RIP-immunoreactive) nerve fiber bundles (Figure 1A) and GFAP-poor tracks (Figure 1B) were observed in the m2cd region. In the GFAP-poor tracks, densely packed myelinated nerve fiber bundles were located (Figure 1A, B). Weak GS-immunoreactivity was detectable in the m2cd region (Figure 2B). Moderately packed myelinated nerve fiber bundles were seen in the m1ab region (Figure 1A). Columns of GFAP-immunoreactive cells were observed to interdigitate with these myelinated nerve fiber bundles (Figure 1A, B). GFAP-rich tracks were present in the m1ab region (Figure 1B). In the GFAP-rich tracks, moderately packed myelinated nerve fiber bundles were located (Figure 1A, B). Moderate GS-immunoreactivity was clearly seen in the m1ab region (Figure 2B). The chemoarchitecture of GFAP, GS, and RIP in the m2ab region resembled that in the m2cd region. Regarding the chemoarchitecture of GFAP, GS, and of RIP, the m1cd region was identified as the transitional area between the m1ab and m2ab regions (Figures 1–2).

### 3.3 GFAP and GFAP-immunoreactive filaments

The m1ab region was the major site of GFAP immunoreactivity localization and the major distribution area of GFAP-immunoreactive filaments in the myelinated region of the normal rat optic nerve (Figures 1, 3, 7).

**FIGURE 7.**
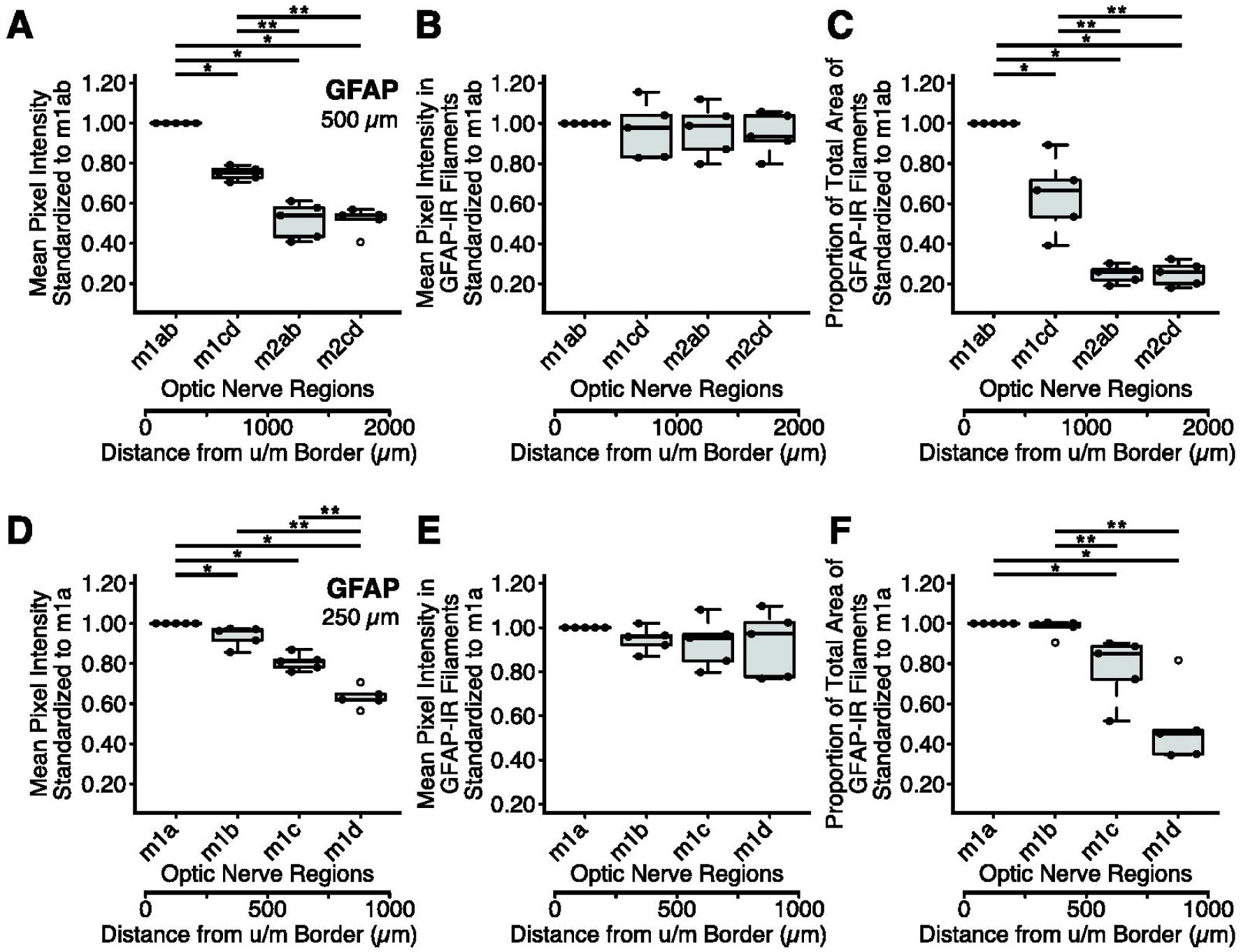
Quantitative analysis of glial fibrillary acidic protein (GFAP) distribution along the course of the myelinated regions in the normal rat optic nerve. **A-B** and **D-E**: The mean standardized pixel intensity within each myelinated region of the optic nerve for GFAP immunoreactivity in each region 500 µm in longitudinal length (**A**) and 250 µm in longitudinal length (**D**), and for GFAP immunoreactivity in GFAP-immunoreactive filaments of each region 500 µm in longitudinal length (**B**) and 250 µm in longitudinal length (**E**). Mean pixel intensity values in regions along the nerve were normalized and expressed as a ratio of the mean pixel intensity of the myelinated region 1ab (m1ab) value (500 µm) or of the myelinated region 1a (m1a) value (250 µm). **C** and **F**: The standardized proportion of the total area of GFAP-immunoreactive filaments within each myelinated region, 500 µm in longitudinal length (**C**) and 250 µm in longitudinal length (**F**). Proportion values in regions along the nerve were normalized and expressed as a ratio of the proportion of the m1ab value (500 µm) or of the m1a value (250 µm). Solid circles on the lines of box and whisker plots indicate the intragroup viability for the chemoarchitectural or structural quantitative measures. The upper whisker shows the maximum value, while the lower whisker represents the minimum value. The thick horizontal line in the box indicates the median. The upper and lower borders of the box show the interquartile range. Open circles represent outliers. The scale at the bottom of each panel indicates the distance from the border between the unmyelinated and myelinated regions (Distance from u/m Border). Note that the intensity of GFAP immunoreactivity was strong in the m1ab region, moderate in myelinated region 1cd (m1cd), and weak in myelinated regions 2ab (m2ab) and 2cd (m2cd). Statistical analyses revealed significant differences in intensity between the m1ab and m1cd regions, and between the m1cd and m2ab regions. However, no significant difference was observed in intensity between the m2ab and m2cd regions (**A**). In myelinated region 1 (m1), GFAP immunoreactivity was the most intense in the m1a region with a gradual decrease in intensity observed along the proximal (posterior) course of the optic nerve. Statistical analyses revealed significant differences in intensity between myelinated regions 1a (m1a) and 1b (m1b), as well as between myelinated regions 1c (m1c) and 1d (m1d). However, there was no significant difference in intensity between the m1b and m1c regions (**D**). *Significant difference with P=0.027 < 0.05, as determined by the Kruskal-Wallis test followed by the Steel-Dwass test for multiple comparisons. **Significant difference with P=0.045 < 0.05, based on the aforementioned statistical procedures.

The intensity of GFAP immunoreactivity was strong in the m1ab region, moderate in the m1cd region, and weak in both the m2ab and m2cd regions. Statistical analyses revealed significant differences in intensity between the m1ab and m1cd regions, as well as between the m1cd and m2ab regions (the Kruskal-Wallis test followed by the Steel-Dwass test for multiple comparisons, P=0.027<0.05 or P=0.045<0.05, n=5). However, no significant difference in intensity was observed between the m2ab and m2cd regions (Figure 7A). In the m1 region, the intensity of GFAP immunoreactivity was the strongest in the m1a region, with a gradual decrease in intensity along the proximal course of the optic nerve. Statistical analyses revealed significant differences in intensity between the m1a and m1b regions, and between the m1c and m1d regions (the Kruskal-Wallis test followed by the Steel-Dwass test for multiple comparisons, P=0.027<0.05 or P=0.045<0.05, n=5). However, no significant difference in intensity was noted between the m1b and m1c regions (Figure 7D).

The proportion of the total area of GFAP-immunoreactive filaments was high in the m1ab region, moderate in the m1cd region, and low in both the m2ab and m2cd regions. Statistical analyses revealed significant differences in the proportion between the m1ab and m1cd regions, as well as between the m1cd and m2ab regions (the Kruskal-Wallis test followed by the Steel-Dwass test for multiple comparisons, P=0.027<0.05 or P=0.045<0.05, n=5). However, there was no significant difference in the proportion between the m2ab and m2cd regions (Figure 7C). In the m1 region, the proportion of the total area of GFAP-immunoreactive filaments was higher in both the m1a and m1b regions, but lower in both the m1c and m1d regions. Statistical analyses revealed significant differences in the proportion between the m1b and m1c regions (the Kruskal-Wallis test followed by the Steel-Dwass test for multiple comparisons, P=0.027<0.05 or P=0.045<0.05, n=5). However, there was no significant difference in the proportion between the m1a and m1b regions, as well as between the m1c and m1d regions (Figure 7F).

The intensity of GFAP immunoreactivity in GFAP-immunoreactive filaments remained largely constant from the m1ab region through the m2cd region. Statistical analyses indicated no significant difference in intensity among the m1ab, m1cd, m2ab, and m2cd regions (the Kruskal-Wallis test followed by the Steel-Dwass test for multiple comparisons, P≧0.05, n=5; Figure 7B). In the m1 region, the intensity of GFAP immunoreactivity in GFAP-immunoreactive filaments also remained largely constant from the m1a region through the m1d region. Statistical analyses indicated no significant difference in intensity among the m1a, m1b, m1c, and m1d regions (the Kruskal-Wallis test followed by the Steel-Dwass test for multiple comparisons, P≧0.05, n=5; Figure 7E).

### 3.4 GS and GS-immunoreactive cells

The m1ab region was the major site of GS immunoreactivity localization in the myelinated region of the normal rat optic nerve (Figures 2, 4, 8).

**FIGURE 8.**
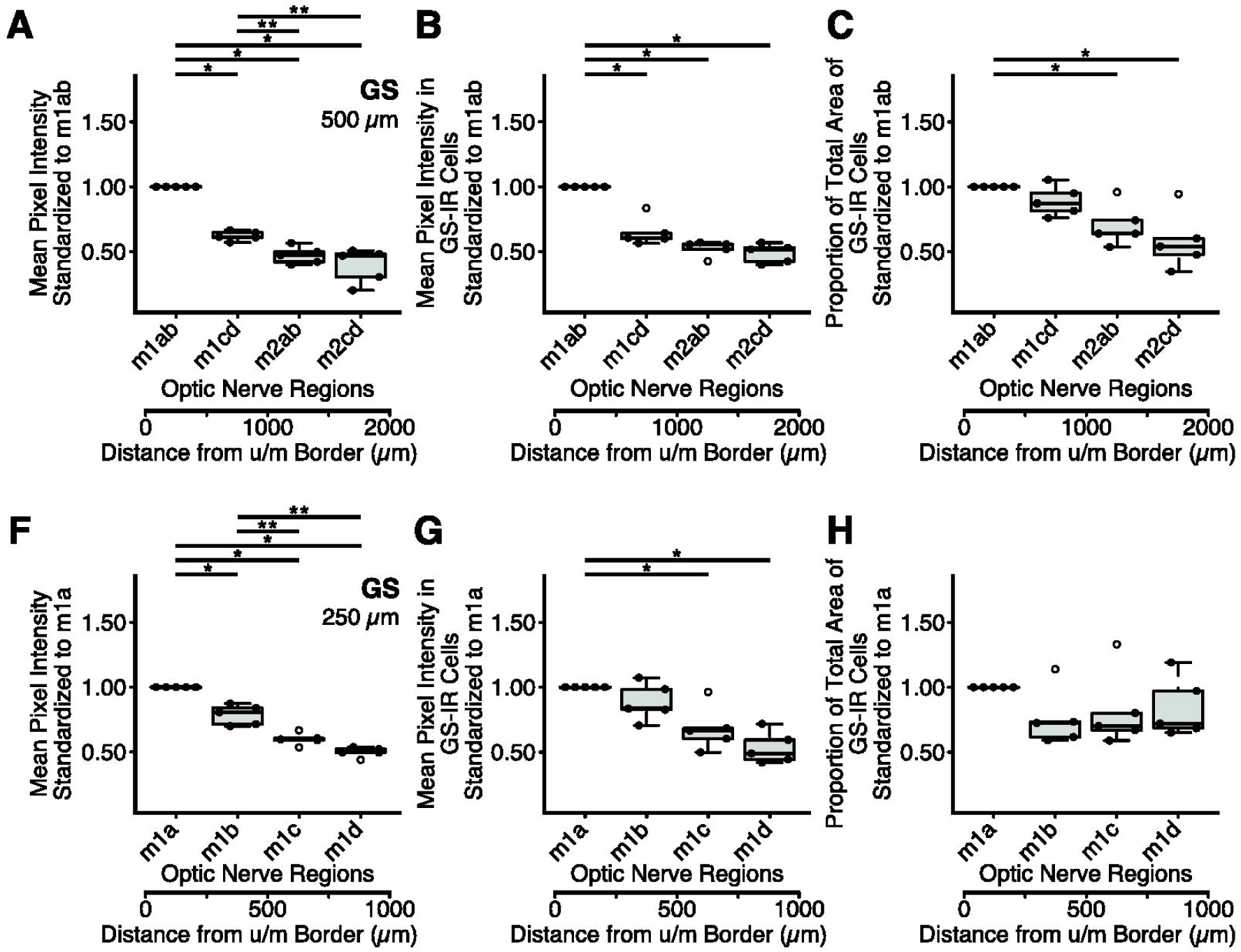
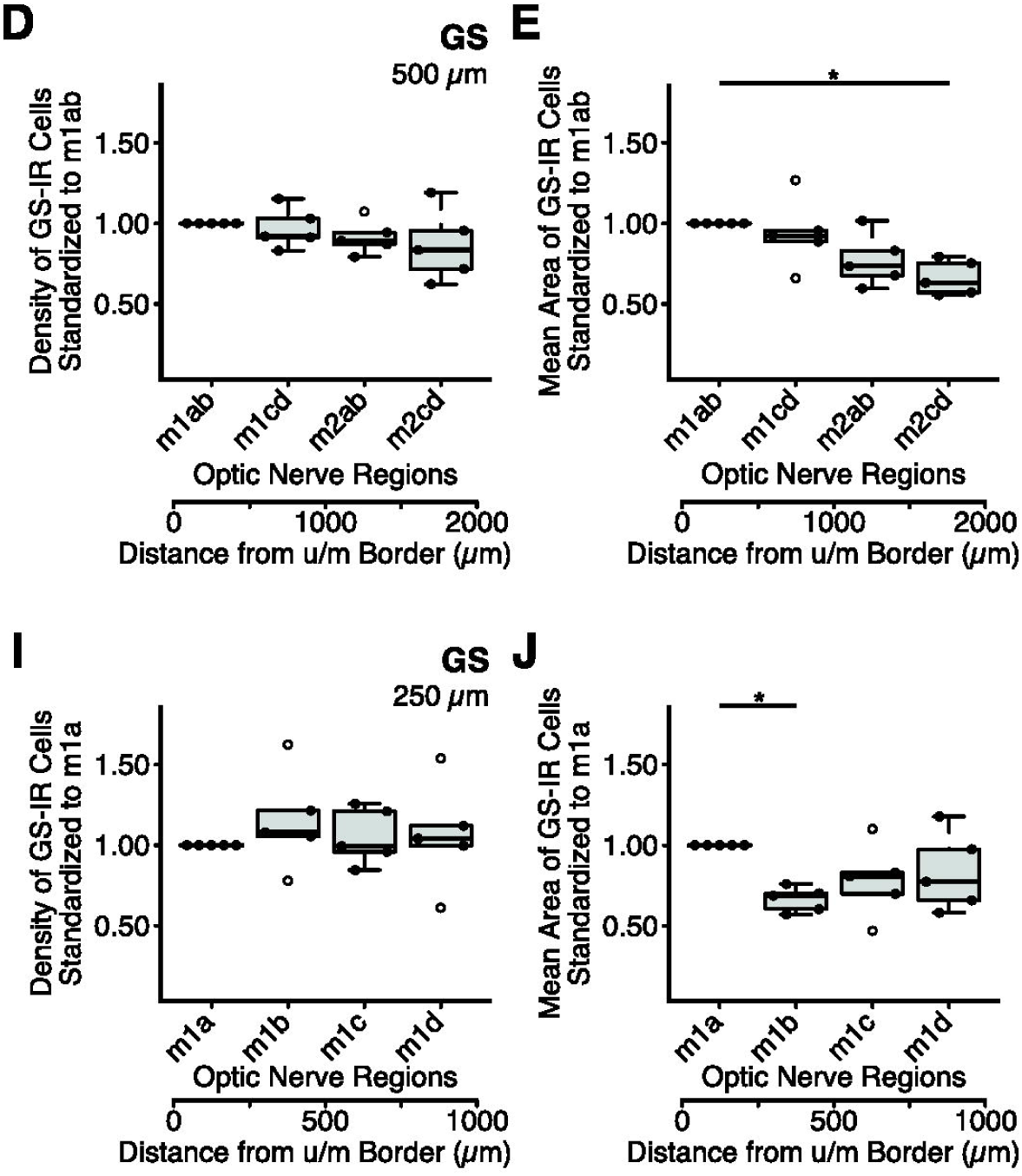
Quantitative analysis of glutamine synthetase (GS) distribution along the course of the myelinated regions in the normal rat optic nerve. **A-B** and **F-G**: The mean standardized pixel intensity within each myelinated region of the optic nerve for GS in each region 500 µm in longitudinal length (**A**) and 250 µm in longitudinal length (**F**), and for GS in GS-immunoreactive cells of each region 500 µm in longitudinal length (**B**) and 250 µm in longitudinal length (**G**). Mean pixel intensity values in regions along the nerve were normalized and expressed as a ratio of the mean pixel intensity of the myelinated region 1ab (m1ab) value (500 µm) or of the myelinated region 1a (m1a) value (250 µm). **C** and **H**: The standardized proportion of the total area of GS-immunoreactive cells within each myelinated region 500 µm in longitudinal length (**C**) and 250 µm in longitudinal length (**H**). Proportion values in regions along the nerve were normalized and expressed as a ratio of the proportion of the m1ab value (500 µm) or of the m1a value (250 µm). **D** and **I**: The standardized density of GS-immunoreactive cells within each myelinated region, 500 µm in longitudinal length (**D**) and 250 µm in longitudinal length (**I**). Density values in regions along the nerve were normalized and expressed as a ratio of the density of the m1ab value (500 µm) or of the m1a value (250 µm). **E** and **J**: The mean standardized area of GS-immunoreactive cells within each myelinated region, 500 µm in longitudinal length (**E**) and 250 µm in longitudinal length (**J**). Mean area values in regions along the nerve were normalized and expressed as a ratio of the mean area of the m1ab value (500 µm) or of the m1a value (250 µm). Solid circles on the lines of box and whisker plots indicate the intragroup viability for the chemoarchitectural or structural quantitative measures. The upper whisker shows the maximum value, while the lower whisker represents the minimum value. The thick horizontal line in the box indicates the median. The upper and lower borders of the box show the interquartile range. Open circles represent outliers. The scale at the bottom of each panel indicates the distance from the border between the unmyelinated and myelinated regions (Distance from u/m Border). Note that the intensity of GS immunoreactivity was strong in the m1ab region, moderate in the myelinated region 1cd (m1cd), and weak in the myelinated regions 2ab (m2ab) and 2cd (m2cd). Statistical analyses revealed significant differences in intensity between the m1ab and m1cd regions, as well as between the m1cd and m2ab regions. However, no significant difference was observed in intensity between the m2ab and m2cd regions (**A**). In myelinated region 1 (m1), GS immunoreactivity was the most intense in the m1a region, moderate in myelinated region 1b (m1b), and the least intense in myelinated regions 1c (m1c) and 1d (m1d). Statistical analyses revealed significant differences in intensity between the m1a and m1b regions, and between the m1b and m1c regions. However, there was no significant difference in intensity between the m1c and m1d regions (**F**). *Significant difference with P=0.027 < 0.05, as determined by the Kruskal-Wallis test followed by the Steel-Dwass test for multiple comparisons. **Significant difference with P=0.045 < 0.05, based on the aforementioned statistical procedures. (The author would like to have the two panels (panels **A-C** (**F-H**) and **D-E** (**I-J**) of Figure 8) on opposite pages.)

The intensity of GS immunoreactivity was strong in the m1ab region, moderate in the m1cd region, and weak in both the m2ab and m2cd regions. Statistical analyses revealed significant differences in intensity between the m1ab and m1cd regions, as well as between the m1cd and m2ab regions (the Kruskal-Wallis test followed by the Steel-Dwass test for multiple comparisons, P=0.027<0.05 or P=0.045<0.05, n=5). However, there was no significant difference in intensity between the m2ab and m2cd regions (Figure 8A). In the m1 region, GS immunoreactivity was most intense in the m1a region, moderate in the m1b region, and least intense in both the m1c and m1d regions. Statistical analyses revealed significant differences in intensity between the m1a and m1b regions, and between the m1b and m1c regions (the Kruskal-Wallis test followed by the Steel-Dwass test for multiple comparisons, P=0.027<0.05 or P=0.045<0.05, n=5). However, there was no significant difference in intensity between the m1c and m1d regions (Figure 8F).

The intensity of GS immunoreactivity in GS-immunoreactive cells was strong in the m1ab region, whereas it was moderate in the remaining three regions: the m1cd, m2ab, and m2cd regions. Statistical analyses revealed significant differences in intensity between the m1ab and m1cd regions, between the m1ab and m2ab regions, and between the m1ab and m2cd regions (the Kruskal-Wallis test followed by the Steel-Dwass test for multiple comparisons, P=0.027<0.05, n=5). However, there was no significant difference in intensity among the m1cd, m2ab, and m2cd regions (Figure 8B). In the m1 region, the intensity of GS immunoreactivity in GS-immunoreactive cells was strong in the m1a region, strong-to-moderate in the m1b region, and moderate in both the m1c and m1d regions. Statistical analyses revealed significant differences in intensity between the m1a and m1c regions, as well as between the m1a and m1d regions (the Kruskal-Wallis test followed by the Steel-Dwass test for multiple comparisons, P=0.027<0.05, n=5). However, there was no significant difference in intensity between the m1a and m1b regions, between the m1b and m1c regions, and between the m1c and m1d regions (Figure 8G).

The proportion of the total area of GS-immunoreactive cells was large in the m1ab region, large-to-medium in the m1cd region, and medium in the m2ab and m2cd regions. Statistical analyses revealed significant differences in the proportion between the m1ab and m2ab regions, and between the m1ab and m2cd regions (the Kruskal-Wallis test followed by the Steel-Dwass test for multiple comparisons, P=0.027<0.05, n=5). However, there was no significant difference in intensity between the m1ab and m1cd regions, between the m1cd and m2ab regions, and between the m2ab and m2cd regions (Figure 8C). In the m1 region, the proportion of the total area of GS-immunoreactive cells remained largely constant from the m1a region through the m1d region. Statistical analyses revealed no significant difference in the proportion among the m1a, m1b, m1c, and m1d regions (the Kruskal-Wallis test followed by the Steel-Dwass test for multiple comparisons, P≧0.05, n=5; Figure 8H).

The mean area of GS-immunoreactive cells was large in the m1ab region, large-to-medium in the m1cd and m2ab regions, and medium in the m2cd region. Statistical analyses revealed significant differences in the mean area between the m1ab and m2cd regions (the Kruskal-Wallis test followed by the Steel-Dwass test for multiple comparisons, P=0.027<0.05, n=5). However, there was no significant difference in the mean area between the m1ab and m1cd regions, between the m1cd and m2ab regions, and between the m2ab and m2cd regions (Figure 8E). In the m1ab region, the mean area of GS-immunoreactive cells in the m1a region was larger than that in the m1b region. Statistical analyses revealed significant differences in the mean area between the m1a and m1b regions (the Kruskal-Wallis test followed by the Steel-Dwass test for multiple comparisons, P=0.027<0.05, n=5; Figure 8J).

The density of GS-immunoreactive cells remained largely constant from the m1ab region through the m2cd region. Statistical analyses indicated no significant difference in density among the m1ab, m1cd, m2ab, and m2cd regions (the Kruskal-Wallis test followed by the Steel-Dwass test for multiple comparisons, P≧0.05, n=5; Figure 8D). In the m1 region, the density of GS-immunoreactive cells was also largely constant from the m1a region through the m1d region. Statistical analyses indicated no significant difference in density among the m1a, m1b, m1c, and m1d regions (the Kruskal-Wallis test followed by the Steel-Dwass test for multiple comparisons, P≧0.05, n=5; Figure 8I).

Interestingly, the m1a region was the major site of GS distribution in the myelinated region of the normal rat optic nerve; however, GS immunoreactivity in this region was markedly weaker than that in the retina (Figures 2A, G, 8A, F).

### 3.5 bBM and bBM-labeled cell nuclei

The m1ab region was the major site of bBM labeling localization in the myelinated region of the normal rat optic nerve (Figures 2, 5, 9).

**FIGURE 9.**
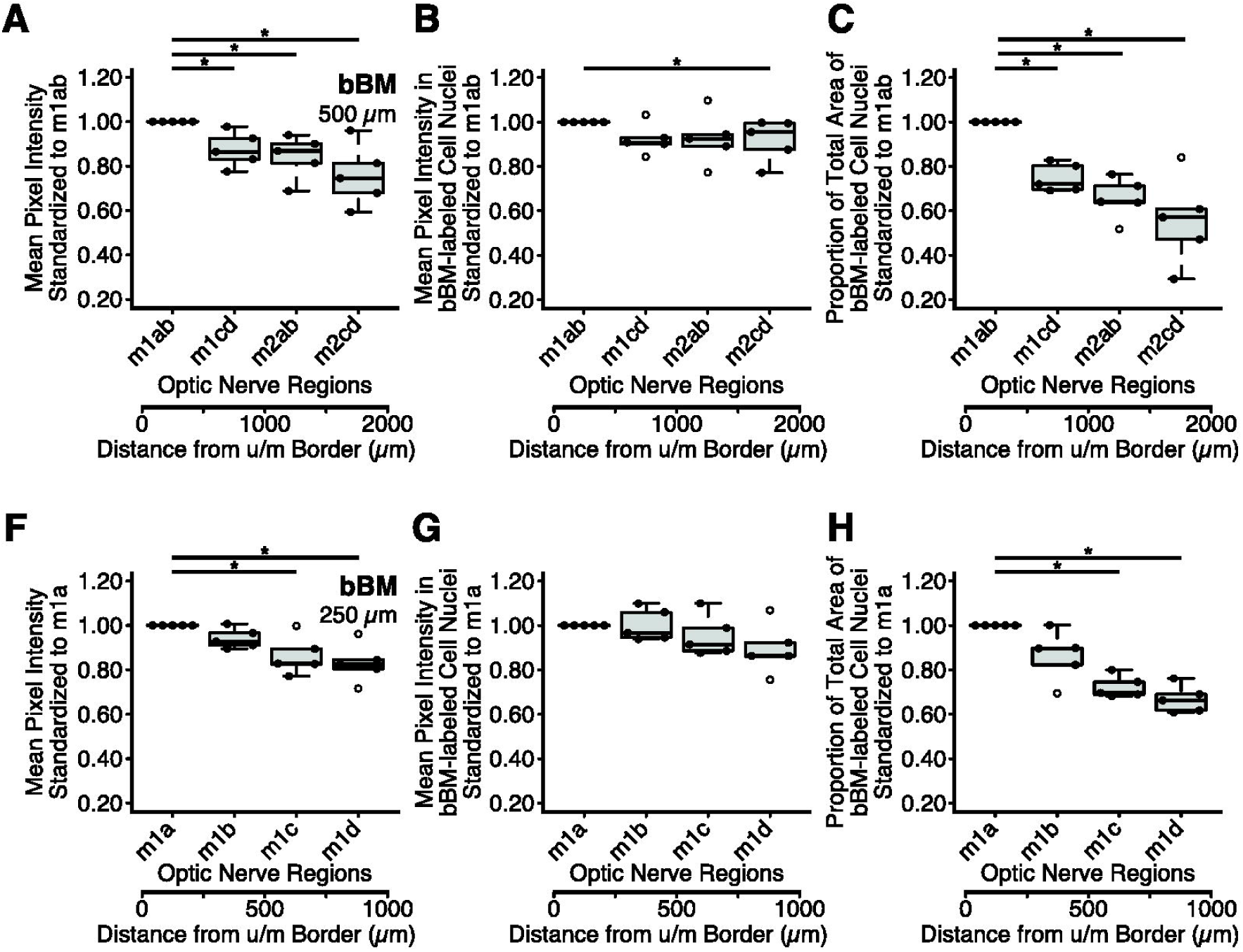
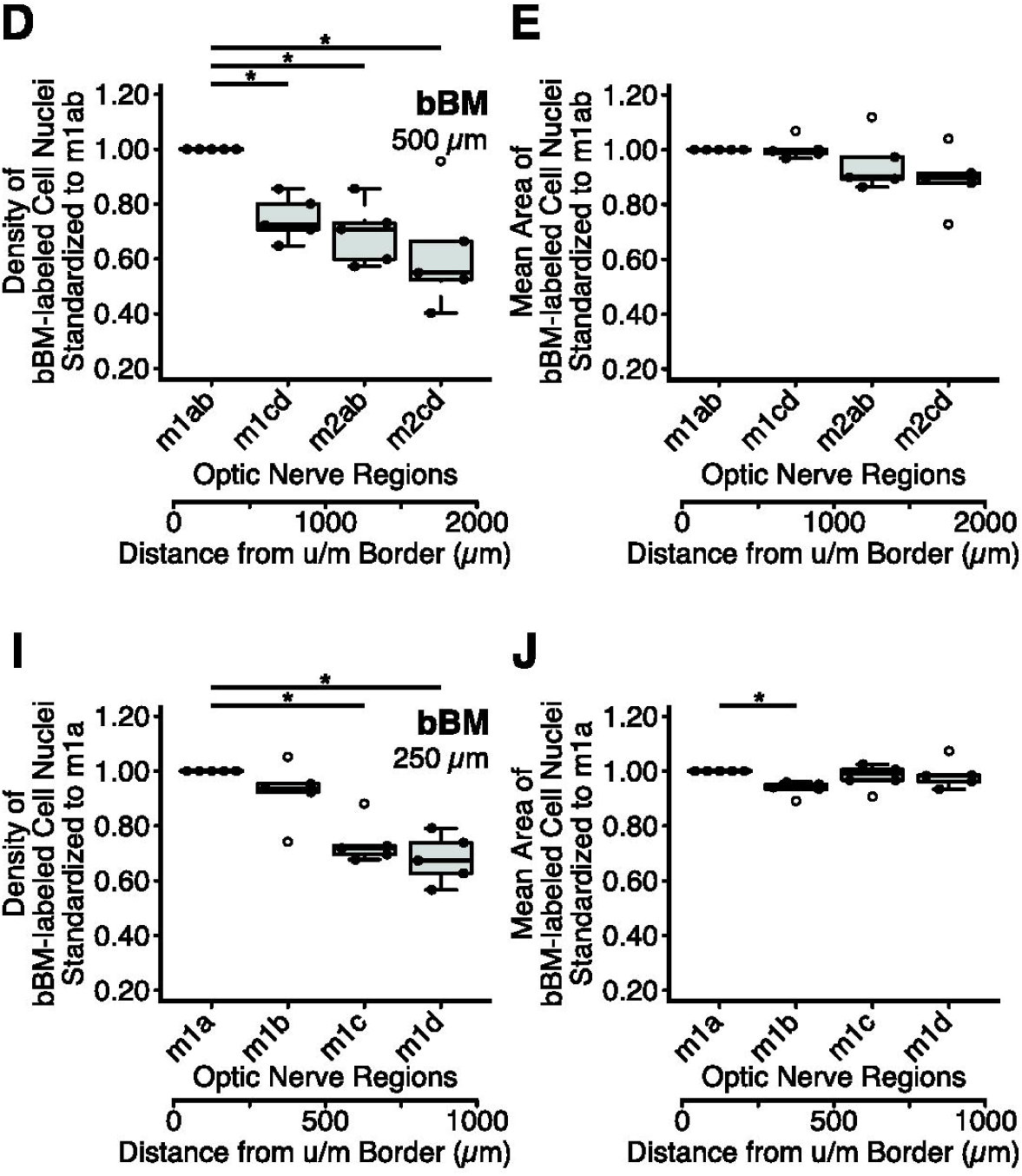
Quantitative analysis of bisBenzimide (bBM; a fluorescent dye for DNA) distribution along the course of the myelinated regions in the normal rat optic nerve. **A-B** and **F-G**: The mean standardized pixel intensity within each myelinated region of the optic nerve for bBM in each region 500 µm in longitudinal length (**A**) and 250 µm in longitudinal length (**F**), and for bBM in bBM-labeled cell nuclei of each region 500 µm in longitudinal length (**B**) and 250 µm in longitudinal length (**G**). Mean pixel intensity values in regions along the nerve were normalized and expressed as a ratio of the mean pixel intensity of the myelinated region 1ab (m1ab) value (500 µm) or of the myelinated region 1a (m1a) value (250 µm). **C** and **H**: The standardized proportion of the total area of bBM-labeled cell nuclei within each myelinated region, 500 µm in longitudinal length (**C**) and 250 µm in longitudinal length (**H**). Proportion values in regions along the nerve were normalized and expressed as a ratio of the proportion of the m1ab value (500 µm) or of the m1a value (250 µm). **D** and **I**: The standardized density of bBM-labeled cell nuclei within each myelinated region, 500 µm in longitudinal length (**D**) and 250 µm in longitudinal length (**I**). Density values in regions along the nerve were normalized and expressed as a ratio of the density of the m1ab value (500 µm) or of the m1a value (250 µm). **E** and **J**: The mean standardized area of bBM-labeled cell nuclei within each myelinated region, 500 µm in longitudinal length (**E**) and 250 µm in longitudinal length (**J**). Mean area values in regions along the nerve were normalized and expressed as a ratio of the mean area of the m1ab value (500 µm) or of the m1a value (250 µm). Solid circles on the lines of box and whisker plots indicate the intragroup viability for the chemoarchitectural or structural quantitative measures. The upper whisker shows the maximum value, while the lower whisker represents the minimum value. The thick horizontal line in the box indicates the median. The upper and lower borders of the box show the interquartile range. Open circles represent outliers. The scale at the bottom of each panel indicates the distance from the border between the unmyelinated and myelinated regions (Distance from u/m Border). Note that the intensity of bBM labeling was strong in the m1ab region, and moderate in myelinated regions 1cd (m1cd), 2ab (m2ab), and 2cd (m2cd). Statistical analyses revealed significant differences in intensity between the m1ab and m1cd regions, between the m1ab and m2ab regions, and between the m1ab and m2cd regions. However, no significant difference was observed among the m1cd, m2ab, and m2cd regions (**A**). In myelinated region 1 (m1), the intensity of bBM labeling was strong in the m1a region, strong-to-moderate in myelinated region 1b (m1b), and moderate in myelinated regions 1c (m1c) and 1d (m1d). Statistical analyses revealed significant differences in intensity between the m1a and m1c regions, and between the m1a and m1d regions. However, there was no significant difference in intensity between the m1a and m1b regions, between the m1b and m1c regions, and between the m1c and m1d regions (**F**). *Significant difference with P=0.027 < 0.05, as determined by the Kruskal-Wallis test followed by the Steel-Dwass test for multiple comparisons. (The author would like to have the two panels (panels **A-C** (**F-H**) and **D-E** (**I-J**) of Figure 9) on opposite pages.)

The intensity of bBM labeling was strong in the m1ab region and moderate in the remaining 3 regions: the m1cd, m2ab, and m2cd regions. Statistical analyses revealed significant differences in intensity between the m1ab and m1cd regions, between the m1ab and m2ab regions, and between the m1ab and m2cd regions (the Kruskal-Wallis test followed by the Steel-Dwass test for multiple comparisons, P=0.027<0.05, n=5). However, there was no significant difference in intensity among the m1cd, m2ab, and m2cd regions (Figure 9A). In the m1 region, the intensity of bBM labeling was strong in the m1a region, strong-to-moderate in the m1b region, and moderate in the m1c and m1d regions. Statistical analyses revealed significant differences in intensity between the m1a and m1c regions, as well as between the m1a and m1d regions (the Kruskal-Wallis test followed by the Steel-Dwass test for multiple comparisons, P=0.027<0.05, n=5). However, there was no significant difference in intensity between the m1a and m1b regions, between the m1b and m1c regions, and between the m1c and m1d regions (Figure 9F).

The intensity of bBM labeling in bBM-labeled cell nuclei was strong in the m1ab region, strong-to-moderate in the m1cd and m2ab regions, and moderate in the m2cd region. Statistical analyses revealed a significant difference in intensity between the m1ab and m2cd regions (the Kruskal-Wallis test followed by the Steel-Dwass test for multiple comparisons, P=0.027<0.05, n=5). However, there was no significant difference in intensity among the m1ab, m1cd, and m2ab regions (Figure 9B). In the m1 region, the intensity of bBM labeling in bBM-labeled cell nuclei remained largely constant from the m1a region through the m1d region. Statistical analyses indicated no significant difference in intensity among the m1a, m1b, m1c, and m1d regions (the Kruskal-Wallis test followed by the Steel-Dwass test for multiple comparisons, P≧0.05, n=5; Figure 9G).

The proportion of the total area of bBM-labeled cell nuclei was large in the m1ab region, and medium in the remaining three regions: the m1cd, m2ab, and m2cd regions. Statistical analyses revealed significant differences in the proportion between the m1ab and m1cd regions, between the m1ab and m2ab regions, and between the m1ab and m2cd regions (the Kruskal-Wallis test followed by the Steel-Dwass test for multiple comparisons, P=0.027<0.05, n=5). However, no significant difference was observed in the proportion among the m1cd, m2ab, and m2cd regions (Figure 9C). In the m1 region, the proportion of the total area of bBM-labeled cell nuclei was large in the m1a region, large-to-medium in the m1b region, and medium in the m1c and m1d regions. Statistical analyses revealed significant differences in the proportion between the m1a and m1c regions, as well as between the m1a and m1d regions (the Kruskal-Wallis test followed by the Steel-Dwass test for multiple comparisons, P=0.027<0.05, n=5). However, there was no significant difference in the proportion between the m1a and m1b regions, between the m1b and m1c regions, and between the m1c and m1d regions (Figure 9H).

The density of bBM-labeled cell nuclei was high in the m1ab region, and medium in the remaining three regions: the m1cd, m2ab, and m2cd regions. Statistical analyses revealed significant differences in the density of bBM-labeled cell nuclei between the m1ab and m1cd regions, between the m1ab and m2ab regions, and between the m1ab and m2cd regions (the Kruskal-Wallis test followed by the Steel-Dwass test for multiple comparisons, P=0.027<0.05, n=5). However, no significant difference was observed in density among the m1cd, m2ab, and m2cd regions (Figure 9D). In the m1 region, the density of bBM-labeled cell nuclei was high in the m1a region, high-to-medium in the m1b region, and medium in the m1c and m1d regions. Statistical analyses revealed significant differences in density between the m1a and m1c regions, and between the m1a and m1d regions (the Kruskal-Wallis test followed by the Steel-Dwass test for multiple comparisons, P=0.027<0.05, n=5). However, there was no significant difference in density between the m1a and m1b regions, between the m1b and m1c regions, and between the m1c and m1d regions (Figure 9I).

The mean area of bBM-labeled cell nuclei remained largely constant from the m1ab region through the m2cd region. Statistical analyses indicated no significant difference in the mean area among the m1ab, m1cd, m2ab, and m2cd regions (the Kruskal-Wallis test followed by the Steel-Dwass test for multiple comparisons, P≧0.05, n=5; Figure 9E). In the m1ab region, the mean area of bBM-labeled cell nuclei in the m1a region was larger than that in the m1b region. Statistical analyses revealed a significant difference in the mean area between the m1a and m1b regions (the Kruskal-Wallis test followed by the Steel-Dwass test for multiple comparisons, P=0.027<0.05, n=5; Figure 9J).

### 3.6 MBPh and MBPh-immunoreactive particles

The m1ab region, especially the m1a region, was the major site of MBPh-immunoreactive-particle localization in the myelinated region of the normal rat optic nerve (Figures 2, 6, 10).

**FIGURE 10.**
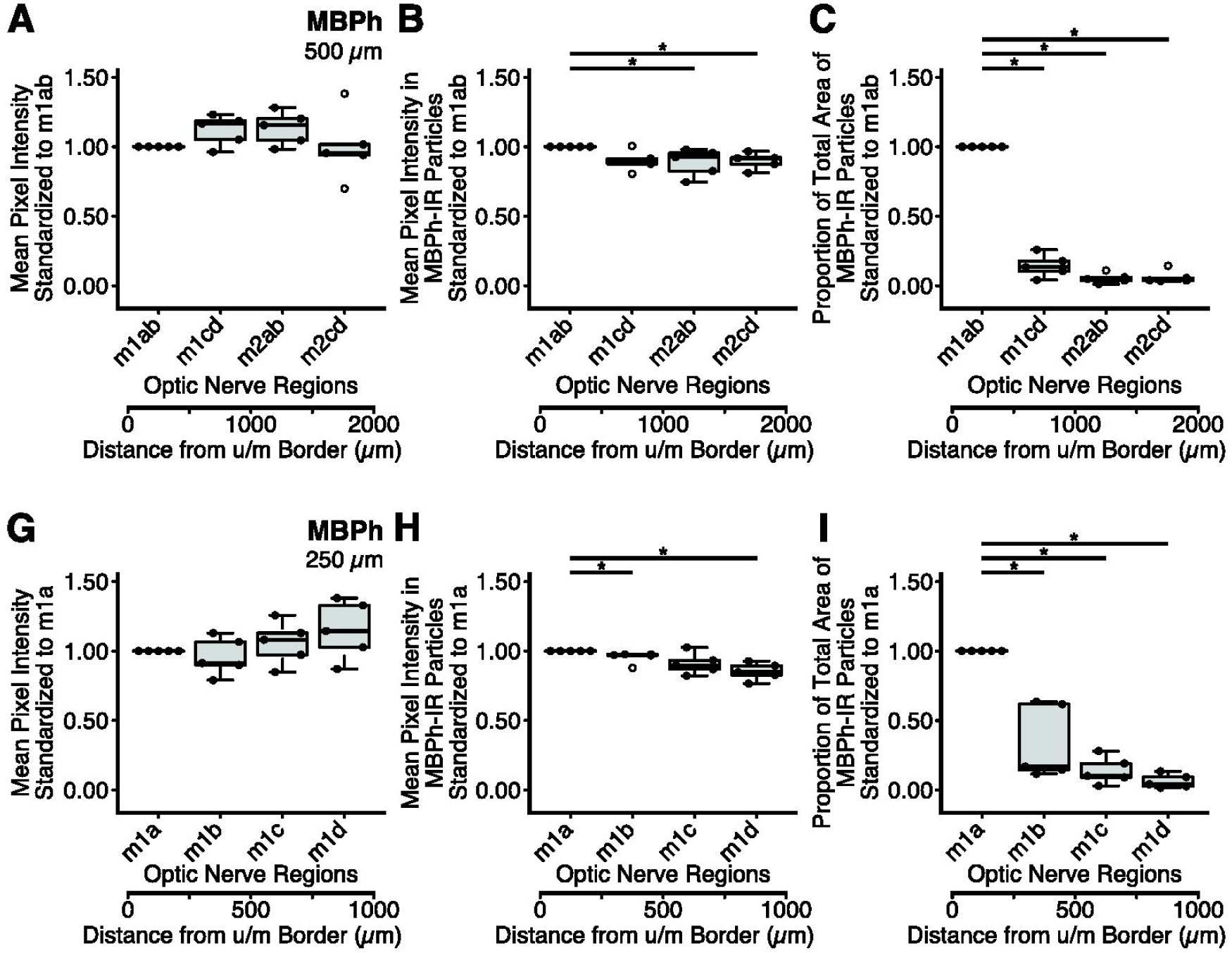
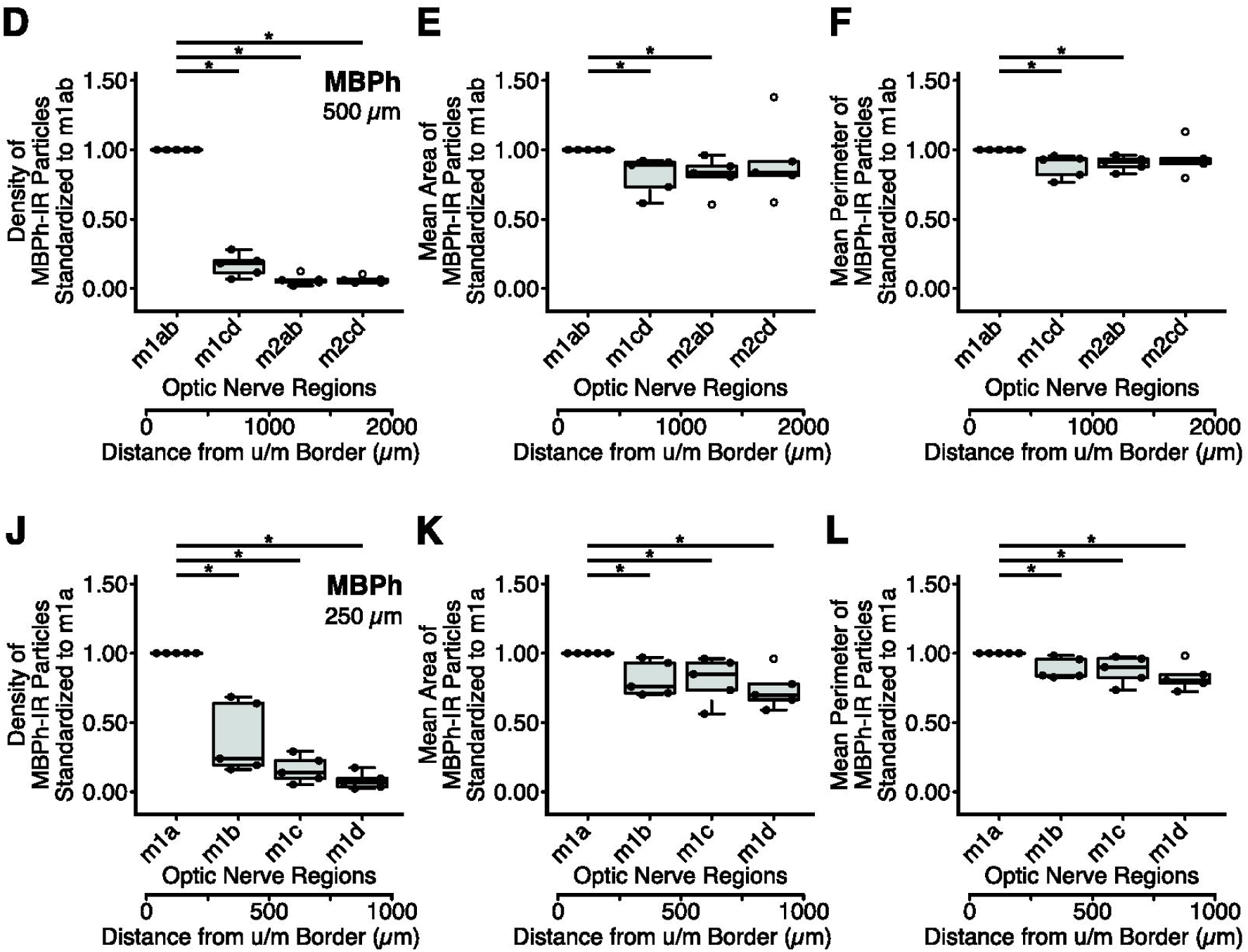
Quantitative analysis of myelin basic protein (MBPh^6^) distribution along the course of the myelinated regions in the normal rat optic nerve. **A-B** and **G-H**: The mean standardized pixel intensity within each myelinated region of the optic nerve for MBPh in each region 500 µm in longitudinal length (**A**) and 250 µm in longitudinal length (**G**), and for MBPh in MBPh-immunoreactive particles of each region 500 µm in longitudinal length (**B**) and 250 µm in longitudinal length (**H**). Mean pixel intensity values in regions along the nerve were normalized and expressed as a ratio of the mean pixel intensity of the myelinated region 1ab (m1ab) value (500 µm) or of the myelinated region 1a (m1a) value (250 µm). **C** and **I**: The standardized proportion of the total area of MBPh-immunoreactive particles within each myelinated region, 500 µm in longitudinal length (**C**) and 250 µm in longitudinal length (**I**). Proportion values in regions along the nerve were normalized and expressed as a ratio of the proportion of the m1ab value (500 µm) or of the m1a value (250 µm). **D** and **J**: The standardized density of MBPh-immunoreactive particles within each myelinated region, 500 µm in longitudinal length (**D**) and 250 µm in longitudinal length (**J**). Density values in regions along the nerve were normalized and expressed as a ratio of the density of the m1ab value (500 µm) or of the m1a value (250 µm). **E** and **K**: The mean standardized area of MBPh-immunoreactive particles within each myelinated region, 500 µm in longitudinal length (**E**) and 250 µm in longitudinal length (**K**). Mean area values in regions along the nerve were normalized and expressed as a ratio of the mean area of the m1ab value (500 µm) or of the m1a value (250 µm). **F** and **L**: The mean standardized perimeter of MBPh-immunoreactive particles within each myelinated region, 500 µm in longitudinal length (**F**) and 250 µm in longitudinal length (**L**). Mean perimeter values in regions along the nerve were normalized and expressed as a ratio of the mean perimeter of the m1ab value (500 µm) or of the m1a value (250 µm). Solid circles on the lines of box and whisker plots indicate the intragroup viability for the chemoarchitectural or structural quantitative measures. The upper whisker shows the maximum value, while the lower whisker represents the minimum value. The thick horizontal line in the box indicates the median. The upper and lower borders of the box show the interquartile range. Open circles represent outliers. The scale at the bottom of each panel indicates the distance from the border between the unmyelinated and myelinated regions (Distance from u/m Border). Note that the intensity of MBPh immunoreactivity in MBPh-immunoreactive particles was strong in the m1ab region, strong-to-moderate in myelinated region 1cd (m1cd), and moderate in myelinated regions 2ab (m2ab) and 2cd (m2cd). Statistical analyses revealed significant differences in intensity between the m1ab and m2ab regions, as well as between the m1ab and m2cd regions. However, no significant difference was observed in intensity between the m1ab and m1cd regions, between the m1cd and m2ab regions, and between the m2ab and m2cd regions (**B**). In myelinated region 1 (m1), the intensity of MBPh immunoreactivity in the MBPh-immunoreactive particles was strong in the m1a region, strong-to-moderate in myelinated region 1c (m1c), and moderate in myelinated regions 1b (m1b) and 1d (m1d). Statistical analyses revealed significant differences in intensity of MBPh immunoreactivity between the m1a and m1b regions, as well as between the m1a and m1d regions. However, there was no significant difference in intensity between the m1a and m1c regions, between the m1b and m1c region, and between the m1c and m1d regions (**H**). The mean area was large in the m1ab region, but small in the m1cd, m2ab, and m2cd regions. Statistical analyses revealed significant differences in the mean area between the m1ab and m1cd regions, and between the m1ab and m2ab regions. However, no significant difference was observed in the mean area between the m1ab and m2cd regions (**E**). In the m1 region, the mean area was large in the m1a region but small in the m1b, m1c, and m1d regions. Statistical analyses revealed significant differences in mean area between the m1a and m1b regions, between m1a and m1c regions, and between the m1a and m1d regions. However, there was no significant difference in mean area among the m1b, m1c, and m1d regions (**K**). *Significant difference with P=0.027 < 0.05, as determined by the Kruskal-Wallis test followed by the Steel-Dwass test for multiple comparisons. (The author would like to have the two panels (panels **A-C** (**G-I**) and **D-F** (**J-L**) of Figure 10) on opposite pages.)

The intensity of MBPh immunoreactivity remained largely constant from the m1ab region through the m2cd region. Statistical analyses indicated no significant difference in intensity among the m1ab, m1cd, m2ab, and m2cd regions (the Kruskal-Wallis test followed by the Steel-Dwass test for multiple comparisons, P≧0.05, n=5; Figure 10A). In the m1 region, the intensity of MBPh immunoreactivity also remained largely constant from the m1a region through the m1d region. Statistical analyses indicated no significant difference in intensity among the m1a, m1b, m1c, and m1d regions (the Kruskal-Wallis test followed by the Steel-Dwass test for multiple comparisons, P≧0.05, n=5; Figure 10G).

The intensity of MBPh immunoreactivity in MBPh-immunoreactive particles was strong in the m1ab region, strong-to-moderate in the m1cd region, and moderate in the m2ab and m2cd regions. Statistical analyses revealed significant differences in intensity between the m1ab and m2ab regions, as well as between the m1ab and m2cd regions (the Kruskal-Wallis test followed by the Steel-Dwass test for multiple comparisons, P=0.027<0.05, n=5). However, there was no significant difference in intensity between the m1ab and m1cd regions, between the m1cd and m2ab regions, and between the m2ab and m2cd regions (Figure 10B). In the m1 region, the intensity of MBPh immunoreactivity in MBPh-immunoreactive particles was strong in the m1a region, strong-to-moderate in the m1c region, and moderate in both the m1b and m1d regions. Statistical analyses revealed significant differences in intensity between the m1a and m1b regions, as well as between the m1a and m1d regions (the Kruskal-Wallis test followed by the Steel-Dwass test for multiple comparisons, P=0.027<0.05, n=5). However, no significant difference was observed in intensity between the m1a and m1c regions, between the m1b and m1c regions, and between the m1c and m1d regions (Figure 10H).

The proportion of the total area of MBPh-immunoreactive particles was large in the m1ab region but small in the remaining three regions: the m1cd, m2ab, and m2cd regions. Statistical analyses revealed significant differences in proportion between the m1ab and m1cd regions, between the m1ab and m2ab regions, and between the m1ab and m2cd regions (the Kruskal-Wallis test followed by the Steel-Dwass test for multiple comparisons, P=0.027<0.05, n=5). However, there was no significant difference in proportion among the m1cd, m2ab, and m2cd regions (Figure 10C). In the m1 region, the proportion of the total area of MBPh-immunoreactive particles was large in the m1a region, medium-to-small in the m1b region, and small in both the m1c and m1d regions. Statistical analyses revealed significant differences in proportion between the m1a and m1b regions, between the m1a and m1c regions, and between the m1a and m1d regions (the Kruskal-Wallis test followed by the Steel-Dwass test for multiple comparisons, P=0.027<0.05, n=5). However, no significant difference was observed in proportion among the m1b, m1c, and m1d regions (Figure 10I).

The density of MBPh-immunoreactive particles was high in the m1ab region but low in the remaining three regions: the m1cd, m2ab, and m2cd regions. Statistical analyses revealed significant differences in the density of MBPh-immunoreactive particles between the m1ab and m1cd regions, between the m1ab and m2ab regions, and between the m1ab and m2cd regions (the Kruskal-Wallis test followed by the Steel-Dwass test for multiple comparisons, P=0.027<0.05, n=5). However, there was no significant difference in density among the m1cd, m2ab, and m2cd regions (Figure 10D). In the m1 region, the density of MBPh-immunoreactive particles was high in the m1a region, medium-to-low in the m1b region, and low in both the m1c and m1d regions. Statistical analyses revealed significant differences in density between the m1a and m1b regions, between the m1a and m1c regions, and between the m1a and m1d regions (the Kruskal-Wallis test followed by the Steel-Dwass test for multiple comparisons, P=0.027<0.05, n=5). However, no significant difference was observed in density among the m1b, m1c, and m1d regions (Figure 10J).

The mean area of MBPh-immunoreactive particles was large in the m1ab region, medium in the m1cd and m2ab regions, and large-to-medium in the m2cd region. Statistical analyses revealed significant differences in the mean area between the m1ab and m1cd regions, as well as between the m1ab and m2ab regions (the Kruskal-Wallis test followed by the Steel-Dwass test for multiple comparisons, P=0.027<0.05, n=5). However, there was no significant difference in the mean area between the m1ab and m2cd regions. Additionally, there was no significant difference in the mean area among the m1cd, m2ab, and m2cd regions (Figure 10E). In the m1 region, the mean area of MBPh-immunoreactive particles was large in the m1a region, and medium in the remaining three regions: m1b, m1c, and m1d regions. Statistical analyses revealed significant differences in the mean area between the m1a and m1b regions, between the m1a and m1c regions, and between the m1a and m1d regions (the Kruskal-Wallis test followed by the Steel-Dwass test for multiple comparisons, P=0.027<0.05, n=5). However, no significant difference was observed in the mean area among the m1b, m1c, and m1d regions (Figure 10K).

The mean perimeter of MBPh-immunoreactive particles was long in the m1ab region, medium in the m1cd and m2ab regions, and long-to-medium in the m2cd region. Statistical analyses revealed significant differences in the mean perimeter between the m1ab and m1cd regions, as well as between the m1ab and m2ab regions (the Kruskal-Wallis test followed by the Steel-Dwass test for multiple comparisons, P=0.027<0.05, n=5). However, there was no significant difference in the mean perimeter between the m1ab and m2cd regions. Additionally, there was no significant difference in the mean perimeter among the m1cd, m2ab, and m2cd regions (Figure 10F). In the m1 region, the mean perimeter of MBPh-immunoreactive particles was long in the m1a region and medium in the remaining three regions: the m1b, m1c, and m1d regions. Statistical analyses revealed significant differences in the mean perimeter between the m1a and m1b regions, between the m1a and m1c regions, and between the m1a and m1d regions (the Kruskal-Wallis test followed by the Steel-Dwass test for multiple comparisons, P=0.027<0.05, n=5). However, no significant difference was observed in the mean perimeter among the m1b, m1c, and m1d regions (Figure 10L).

Interestingly, MBPh-immunoreactive particles were densely distributed in the m1ab region, notably in the m1a region; however, these particles were sparsely dispersed in the m1cd, m2ab, and in the m2cd regions (Figure 10).

### 3.7 Shape descriptors of MBPh-immunoreactive particles

The shape descriptors, such as circularity, AR (aspect ratio), roundness, and solidity, of MBPh-immunoreactive particles remained primarily stable from the m1ab region through the m2cd region and from the m1a region through the m1d region, except for the circularity in the m1 region (Figure 11).

**FIGURE 11.**
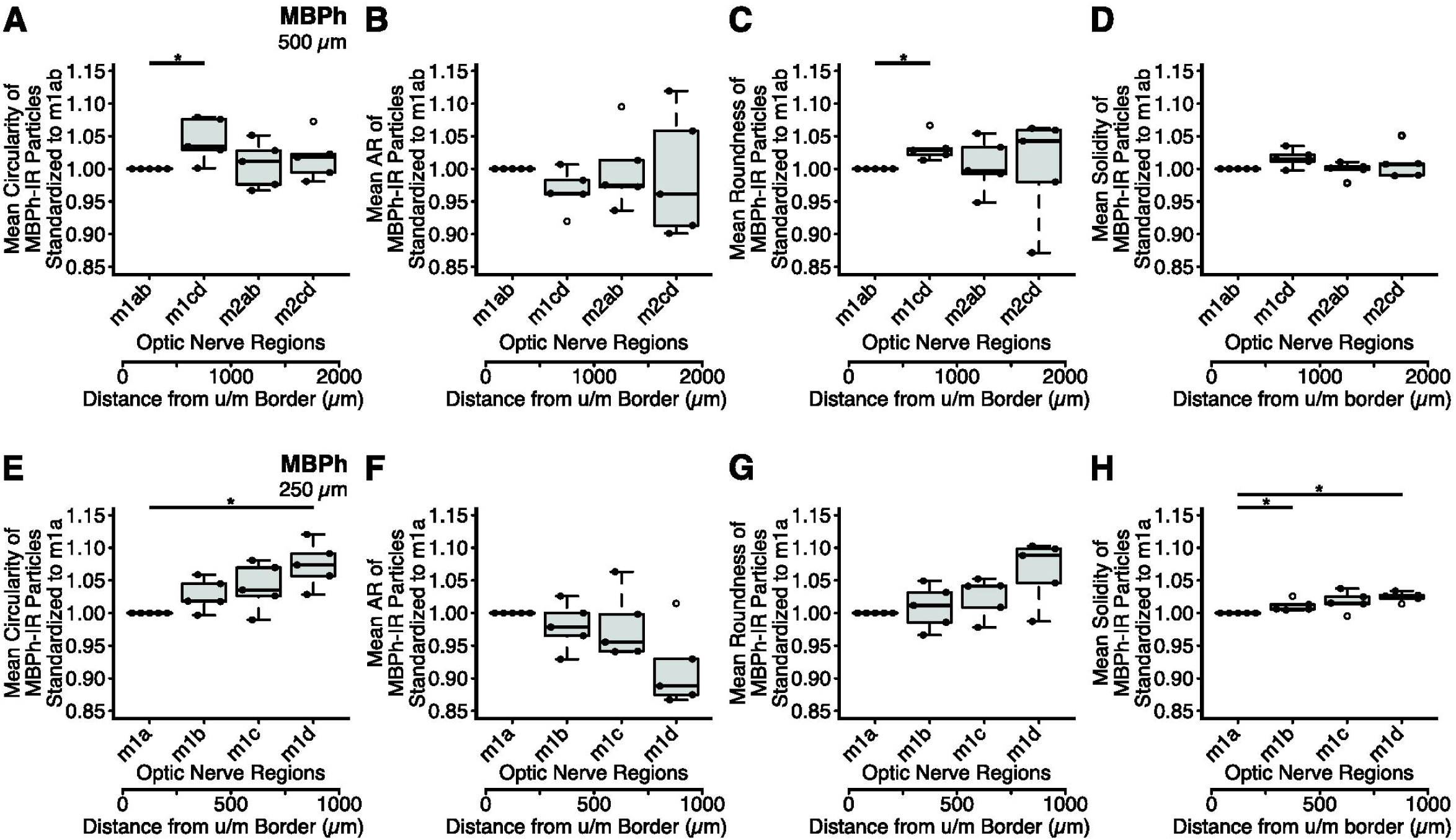
Quantitative analysis of the shape descriptors of myelin basic protein (MBPh^7^)-immunoreactive particles along the course of the myelinated regions in the normal rat optic nerve. **A** and **E**: The mean standardized circularity of MBPh-immunoreactive particles within each myelinated region 500 µm in longitudinal length (**A**) and 250 µm in longitudinal length (**E**). The mean circularity values in regions along the nerve were normalized and expressed as a ratio of the mean circularity of the myelinated region 1ab (m1ab) value (500 µm) or of the myelinated region 1a (m1a) value (250 µm). **B** and **F**: The mean standardized AR (aspect ratio) of MBPh-immunoreactive particles within each myelinated region 500 µm in longitudinal length (**B**) and 250 µm in longitudinal length (**F**). The mean AR values in regions along the nerve were normalized and expressed as a ratio of the mean AR of the m1ab value (500 µm) or of the m1a value (250 µm). **C** and **G**: The mean standardized roundness of MBPh-immunoreactive particles within each myelinated region 500 µm in longitudinal length (**C**) and 250 µm in longitudinal length (**G**). The mean roundness values in regions along the nerve were normalized and expressed as a ratio of the mean roundness of the m1ab value (500 µm) or of the m1a value (250 µm). **D** and **H**: The mean standardized solidity of MBPh-immunoreactive particles within each myelinated region 500 µm in longitudinal length (**D**) and 250 µm in longitudinal length (**H**). The mean solidity values in regions along the nerve were normalized and expressed as a ratio of the mean solidity of the m1ab value (500 µm) or of the m1a value (250 µm). Solid circles on the lines of box and whisker plots indicate the intragroup viability for the structural quantitative measures. The upper whisker shows the maximum value, while the lower whisker represents the minimum value. The thick horizontal line in the box indicates the median. The upper and lower borders of the box show the interquartile range. Open circles represent outliers. The scale at the bottom of each panel indicates the distance from the border between the unmyelinated and myelinated regions (Distance from u/m Border). Note that the mean circularity was medium in the m1ab region, large in myelinated region 1cd (m1cd), and small-to-large in myelinated regions 2ab (m2ab) and 2cd (m2cd). Statistical analyses revealed a significant difference in the mean circularity between the m1ab and m1cd regions. However, no significant difference was observed in the mean circularity among the m1cd, m2ab, and m2cd regions (**A**). In myelinated region 1 (m1), the mean circularity was medium in the m1a region, medium-to-large in myelinated regions 1b (m1b) and 1c (m1c), and large in myelinated region 1d (m1d). Statistical analyses revealed a significant difference in the mean circularity between the m1a and m1d regions. However, there was no significant difference in the mean circularity between the m1a and m1b regions, between the m1b and m1c regions, and between the m1c and m1d regions (**E**). *Significant difference with P=0.027 < 0.05, as determined by the Kruskal-Wallis test followed by the Steel-Dwass test for multiple comparisons.

The mean circularity was medium in the m1ab region, large in the m1cd region, and small-to-large in both the m2ab and m2cd regions. Statistical analyses revealed a significant difference in the mean circularity between the m1ab and m1cd regions (the Kruskal-Wallis test followed by the Steel-Dwass test for multiple comparisons, P=0.027<0.05, n=5). However, there was no significant difference in the mean circularity among the m1cd, m2ab, and m2cd regions (Figure 11A). In the m1 region, the mean circularity was medium in the m1a region, medium-to-large in the m1b and m1c regions, and large in the m1d region. Statistical analyses revealed a significant difference in the mean circularity between the m1a and m1d regions (the Kruskal-Wallis test followed by the Steel-Dwass test for multiple comparisons, P=0.027<0.05, n=5). However, no significant difference was observed in the mean circularity between the m1a and m1b regions, between the m1b and m1c regions, and between the m1c and m1d regions (Figure 11E).

The mean AR of MBPh-immunoreactive particles remained largely constant from the m1ab region through the m2cd region. Statistical analyses indicated no significant difference in the mean AR among the m1ab, m1cd, m2ab, and m2cd regions (the Kruskal-Wallis test followed by the Steel-Dwass test for multiple comparisons, P≧0.05, n=5; Figure 11B). In the m1 region, the mean AR of MBPh-immunoreactive particles also remained largely constant from the m1a region through the m1d region. Statistical analyses indicated no significant difference in the mean AR among the m1a, m1b, m1c, and m1d regions (the Kruskal-Wallis test followed by the Steel-Dwass test for multiple comparisons, P≧0.05, n=5; Figure 11F).

The mean roundness was medium in the m1ab region, large in the m1cd region, and small-to-large in both the m2ab and m2cd regions. Statistical analyses revealed a significant difference in the mean roundness between the m1ab and m1cd regions (the Kruskal-Wallis test followed by the Steel-Dwass test for multiple comparisons, P=0.027<0.05, n=5). However, there was no significant difference in the mean circularity among the m1cd, m2ab, and m2cd regions (Figure 11C). In the m1 region, the mean roundness remained largely constant from the m1a region through the m1d region. Statistical analyses indicated no significant difference in the mean roundness among the m1a, m1b, m1c, and m1d regions (the Kruskal-Wallis test followed by the Steel-Dwass test for multiple comparisons, P≧0.05, n=5; Figure 11G).

The mean solidity of MBPh-immunoreactive particles remained largely constant from the m1ab region through the m2cd region. Statistical analyses indicated no significant difference in the mean solidity among the m1ab, m1cd, m2ab, and m2cd regions (the Kruskal-Wallis test followed by the Steel-Dwass test for multiple comparisons, P≧0.05, n=5; Figure 11D). In the m1 region, the mean solidity of MBPh-immunoreactive particles was medium in the m1a region, medium-to-large in the m1c region, and large in both the m1b and m1d regions. Statistical analyses revealed a significant difference in the mean solidity between the m1a and m1b regions, as well as between the m1a and m1d regions (the Kruskal-Wallis test followed by the Steel-Dwass test for multiple comparisons, P=0.027<0.05, n=5). However, there was no significant difference in the mean solidity between the m1a and m1c regions. Additionally, there was no significant difference in the mean solidity among the m1b, m1c, and m1d regions (Figure 11H).

## 4 Discussion

### 4.1 A summary of the results

We examined the distribution patterns of glial cell marker proteins (GFAP, MBPh, and GS) and that of the cell nucleus marker (bBM) in the myelinated region of the normal rat optic nerve. Additionally, we analyzed the distribution pattern and shape descriptors of MBPh-immunoreactive particles in this region. GFAP immunoreactivity and GS immunoreactivity were significantly strong in the m1ab region, moderate in the m1cd region, and weak in both the m2ab and m2cd regions. Similar to the distribution patterns of GFAP immunoreactivity and of GS immunoreactivity, bBM labeling was significantly strong in the m1ab region and moderate in the m1cd region; however, bBM labeling was also moderate in both the m2ab and m2cd regions. In contrast to these findings, MBPh immunoreactivity remained largely constant from the m1ab region through the m2cd region. Interestingly, MBPh-immunoreactive particles were densely distributed in the m1ab region, notably in the m1a region, while these particles were loosely dispersed in the m1cd, m2ab, and m2cd regions. The shape descriptors of these particles remained primarily stable from the m1ab region through the m2cd region, except for the circularity in the m1 region.

### 4.2 Comparison with previous findings

The division scheme used in this study was largely consistent with that of a previous study in the rat, except for the subregions in the myelinated region (Kawano 2015). In both studies, the myelinated region was divided into two subregions: the m1 and m2 regions. In the previous study, the border between the m1*^2^ and m2* regions was defined by the shape of the outline in the myelinated region. The longitudinal length of the m1* region was 872.2 ± 30.9 µm (Average ± SE; n = 5; Kawano 2015). In the present study, the boundary between the m1 and m2 regions was determined at the line where the distance from the u/m border (border between the unmyelinated and myelinated regions) measured 1 mm. Additionally, the proximal (posterior) boundary of the m2 region was set at the line where the distance from the u/m border measured 2 mm. Consequently, the longitudinal length of the m2 region was 1 mm. Therefore, the boundary between the m1 and m2 regions, as well as the proximal (posterior) boundary of the m2 region, were strictly defined. Furthermore, the division scheme used in the present study was systematically organized in the myelinated region.

Kawano (2015) described that GFAP immunoreactivity in the m1* region was stronger than that in the m2* region, with a statistically significant difference. The results obtained in the present study were compatible with these findings. Kawano (2015) also described that GS immunoreactivity in the m1* region was stronger than that in the m2* region; however, no significant difference was observed in the strength of GS immunoreactivity between the m1* and m2* regions. The present study confirmed the distribution pattern of GS immunoreactivity in the m1* and m2* regions, and reported that there was a significant difference in the strength of GS immunoreactivity between the m1ab and m2ab regions, between the m1ab and m2cd regions, between the m1cd and m2ab regions, and between the m1cd and m2cd regions.

Kawano (2026) demonstrated the concentration of MBPh-immunoreactive particles in the distal (anterior)-most part of the myelinated region. The present study confirmed these findings.

### 4.3 Characteristics of the results obtained from the quantitative morphological analyses in each myelinated region

#### 4.3.1 Relationship between the intensity of GFAP immunoreactivity in each myelinated region and the area occupied by GFAP-immunoreactive filaments

The intensity of GFAP immunoreactivity was significantly strong in the m1ab region, moderate in the m1cd region, and weak in both the m2ab and m2cd regions (Figure 7A). This diversity was attributable to significant differences in the proportion of the total area occupied by GFAP-immunoreactive filaments (astrocytic filaments): the proportion was significantly high in the m1ab region, moderate in the m1cd region, and low in both the m2ab and m2cd regions (Figure 7C). This interpretation was supported by the finding that there was no significant difference in intensity of GFAP immunoreactivity in GFAP-immunoreactive filaments among the m1ab, m1cd, m2ab, and m2cd regions (Figure 7B).

#### 4.3.2 Relationship between the intensity of GS immunoreactivity in each myelinated region and that in GS-immunoreactive cells

In the m1 and m2 regions, the intensity of GS immunoreactivity was significantly strong in the m1ab region, moderate in the m1cd region, and weak in both the m2ab and m2cd regions (Figure 8A). These dissimilarities were attributable to significant differences in intensity of GS immunoreactivity in GS-immunoreactive cells: the intensity was strong in the m1ab region and moderate in the m1cd, m2ab, and m2cd regions (Figure 8B). The dissimilarities were also attributed to significant differences in the proportion of the total area occupied by GS-immunoreactive cells: the proportion was large in the m1ab region and medium in both the m2ab and m2cd regions (Figure 8C).

In the m1 region, the intensity of GS immunoreactivity was the strongest in the m1a region, moderate in the m1b region, and the weakest in both the m1c and m1d regions, with statistically significant differences (Figure 8F). This diversity was attributable to significant differences in intensity of GS immunoreactivity in GS-immunoreactive cells: the intensity was significantly strong in the m1a region and moderate in both the m1c and m1d regions (Figure 8G). It is possible that this diversity was also attributed to the proportion of the total area occupied by GS-immunoreactive cells, since four out of five proportions in each region were less than 1.00 from the m1b region through the m1d region, and since the median of the proportion in each region was approximately 0.7 from the m1b region through the m1d region: m1b, 0.72; m1c, 0.70; m1d, 0.72 (Figure 8H).

#### 4.3.3 Relationship between the intensity of bBM-labeling in each myelinated region and that in bBM-labeled cell nuclei

The intensity of bBM-labeling was significantly strong in the m1ab region and moderate in the m1cd, m2ab, and m2cd regions (Figure 9A). This dissimilarity was largely attributable to significant differences in the proportion of the total area occupied by bBM-labeled cell nuclei: the proportion was large in the m1ab region and moderate in the m1cd, m2ab and m2cd regions (Figure 9C). This interpretation was supported by the fact that there was no significant difference in intensity of bBM-labeling in bBM-labeled cell nuclei among the m1ab, m1cd, and m2ab regions (Figure 9B).

Additionally, the diversity in the proportion (of the total area occupied by bBM-labeled cell nuclei) was attributed to significant differences in the density of bBM-labeled cell nuclei: the density was significantly high in the m1ab region and moderate in the m1cd, m2ab, and m2cd regions (Figure 9D). This interpretation was further supported by the fact that there was no significant difference in the mean area of bBM-labeled cell nuclei among the m1ab, m1cd, m2ab, and m2cd regions (Figure 9E).

#### 4.3.4 The intensity of MBPh immunoreactivity in each myelinated region, as well as the measurement values of MBPh-immunoreactive particles, including their shape descriptors

There were significant differences in five measurement values of MBPh-immunoreactive particles between the m1ab region and the remaining three regions: the m1cd, m2ab, and m2cd regions. These five measurement values included the intensity of MBPh immunoreactivity in MBPh-immunoreactive particles, the proportion of the total area occupied by the particles, the density, the mean area, and the mean perimeter of the particles. Among these, the proportion of the total area and the density showed marked differences between the m1ab and the remaining three regions: the m1cd, m2ab, and m2cd regions (Figure 10B-F). The diversity of the proportion was attributable to differences in density. This interpretation was supported by the fact that the differences in the mean area were minor between the m1ab region and the remaining three regions, although there were significant differences in the mean area between the m1ab and m1cd regions, as well as between the m1ab and m2ab regions.

On the other hand, there was no significant difference in intensity of MBPh immunoreactivity among the m1ab, m1cd, m2ab, and m2cd regions (Figure 10A). Therefore, the dissimilarity in the five measurement values of MBPh-immunoreactive particles did not impact the consistency of MBPh immunoreactivity intensity from the m1ab region through the m2cd region.

The shape descriptors of MBPh-immunoreactive particles remained primarily stable from the m1ab region through the m2cd region, except for the circularity and the roundness (Figure 11A-D). The roundness was significantly smaller in the m1ab region than that in the m1cd region (Figure 11C). However, this finding was not supported by the lack of significant difference in roundness among the m1a, m1b, m1c, and m1d regions (Figure 11G). Regarding the circularity in the m1 region, it was significantly smaller in the m1ab region than that in the m1cd region (Figure 11A). The reliability of this finding was strengthened by the fact that circularity was significantly smaller in the m1a region than that in the m1d region (Figure 11E). Therefore, the shape descriptors of MBPh-immunoreactive particles were primarily stable from the m1ab region through the m2cd region, except for the circularity in the m1 region.

### 4.4 Comparisons between the accumulation of GFAP-immunoreactive filaments in the m1ab region and reactive astrogliosis

This section examines the factors that contribute to the accumulation of GFAP-immunoreactive filaments in the m1ab region. In this context, reactive astrogliosis serves as a useful reference for comparing the accumulation. Reactive astrogliosis is a fibrous proliferation of glial cells in injured areas of the central nervous system (CNS; Messam et al. 2002; McAteer and Choudhury 2009; Sofroniew 2009). Almost all brain lesions involve some component of reactive astrogliosis, even in cases of different glial pathologies. Reactive astrogliosis is a secondary response to damage in the CNS and may persist for weeks or months after brain injury. This phenomenon occurs after infarction and is associated with infections and neoplasms, as well as with demyelinating, toxic, and metabolic diseases. During reactive astrogliosis, astrocytes undergo hypertrophy; their nuclei become enlarged, chromatin becomes less dense, and nucleoli become more prominent. There is an increase in the number of organelles and higher production of intermediate filaments such as GFAP, nestin, and vimentin, which results in greater and more highly condensed glial processes and fibers (Messam et al. 2002; McAteer and Choudhury 2009; Sofroniew 2009). Therefore, higher production of GFAP is observed not only in reactive astrogliosis but also in the m1ab region, where GFAP-immunoreactive filaments have accumulated. Since reactive astrogliosis is a secondary response to CNS insults, it is possible that the accumulation of GFAP-immunoreactive filaments in the m1ab region is attributable to physiologically stressed conditions present in this region.

In reactive astrogliosis, the nuclei of astrocytes become enlarged (Messam et al. 2002). In the present study, the mean area of bBM-labeled cell nuclei in the m1ab region was similar to the remaining three myelinated regions: the m1cd, m2ab, and m2cd regions (Figure 9E). Since these cell nuclei were not only parts of astrocytes but also those of oligodendrocytes, microglia, endothelial cells, and of pericytes (Robison et al. 1989; Bron et al. 1997), there is a slight possibility that the mean area of astrocyte nuclei is larger in the m1ab region than in the remaining three regions. In the present study, the density of bBM-labeled cell nuclei was significantly higher in the m1ab region than in the remaining three regions (Figure 9D). There is a distinct possibility that this diversity of the density is attributable to the higher density of astrocyte nuclei in the m1ab region. This interpretation is supported by two facts. One is that the majority (88-98%) of GS-immunoreactive cells are oligodendrocytes in the rat optic nerve (Kawano 2015). Another is that the density of GS-immunoreactive cells in the m1ab region was similar to that in the remaining three regions (Figure 8D). In order to improve our assertion that the m1ab region is under physiologically stressed conditions, further studies are required to clarify both the area and density of astrocyte nuclei in the m1ab region and in the remaining three regions.

### 4.5 The relationship between the strong GS immunoreactivity in oligodendrocytes distributed in the m1ab region and physiologically stressed conditions

GS expression in oligodendrocytes is increased in chronic pathological conditions in the mouse. Under these conditions, the mouse shows a significant increase in the number of GS-immunoreactive cell bodies, as well as a tendency for increased GS immunoreactivity per oligodendrocyte at end-stage compared to control subjects (Ben Haim et al. 2021). In the present study, the density of GS-immunoreactive cells remained largely constant from the m1ab region through the m2cd region. However, a significant increase in GS immunoreactivity was observed in GS-immunoreactive cells of the m1ab region compared to those in the remaining three regions: m1cd, m2ab, and m2cd regions. Accordingly, the significant increase in GS immunoreactivity observed in GS-immunoreactive cells within the m1ab region might be attributable to physiologically stressed conditions rather than physiological ones in this area.

This interpretation is also supported by evidence related to axo-myelinic neurotransmission and oligodendrocyte myelination. In the physiological state, the myelination of axons by oligodendrocytes is a highly complex cell-to-cell interaction. Oligodendrocytes and axons maintain a reciprocal signaling relationship (axo-myelinic neurotransmission), in which oligodendrocytes receive cues from axons that guide their myelination process. Subsequently, oligodendrocytes shape axonal structure and conduction (oligodendrocyte myelination; Duncan et al. 2021). Regarding axo-myelinic neurotransmission in the rat optic nerve, glutamate is a key signaling molecule (Barron and Kim 2019), since retino-geniculate neurons are glutamatergic (Salt 1986; Kielland and Heggelund 2002; Govindaiah et al. 2012). During the propagation of action potentials, the traversing action potentials induce depolarization of the axon. The depolarization is sensed by voltage-gated Ca^2+^ channels, which in turn activate and cause intra-axonal Ca^2+^ release from the axoplasmic reticulum. This process stimulates the fusion of glutamatergic vesicles and the subsequent release of glutamate into the peri-axonal space. The released glutamate then activates myelinic AMPA and NMDA receptors located on the innermost myelin sheaths, finally promoting Ca^2+^ influx into the myelin cytoplasm (Micu et al. 2016; Micu et al. 2018; Barron and Kim 2019; Bergaglio et al. 2021; Duncan et al. 2021). The Ca^2+^ influx is able to lead to various downstream Ca^2+^ pathways important for myelination and oligodendrocyte function (Barron and Kim 2019). Glutamate in the peri-axonal space is then likely taken up by glutamate transporters on axons for reloading into vesicles (Micu et al. 2018), as well as by those on oligodendrocytes for transforming glutamate into glutamine by using GS, since the majority (88-98%) of GS-immunoreactive cells are oligodendrocytes in the rat optic nerve (Kawano 2015).

In the pathological state, excessive vesicular glutamate release into the peri-axonal space is proposed (Micu et al. 2018; Bergaglio et al. 2021). Accordingly, glutamate clearance is required in this space in order to prevent excitotoxicity induced by immoderate glutamate accumulation. To clear excessive glutamate in the peri-axonal space, glutamate is taken up by using glutamate transporters on axons for reloading into vesicles, and by those on oligodendrocytes for transforming glutamate into glutamine through the enzyme glutamine synthetase (GS). Since a large quantity of glutamate is taken into these cells, GS expression is increased in these oligodendrocytes. Therefore, it is possible that the significant increase in GS immunoreactivity observed in GS-immunoreactive cells within the m1ab region is attributable to the physiologically stressed conditions rather than physiological ones in this area.

### 4.6 The relationship between myelin debris-like MBPh-immunoreactive particles distributed in the m1ab region and physiologically stressed conditions

A substantial amount of myelin debris has been documented under pathological conditions (Weinger et al. 2011; Zhou et al. 2019; Wang et al. 2020; Yao et al. 2023). In the present study, the density of MBPh-immunoreactive particles was high in the m1ab region (Figure 10D), especially in the m1a region (Figure 10J). The immunohistochemical characteristics of these MBP-immunoreactive particles were similar to those of myelin debris (Kawano 2026). Therefore, it is probable that the m1ab region was under physiologically stressed conditions, since this region is rich in myelin debris-like MBPh-immunoreactive particles (Figure 10D), and constitutes part of the normal rat optic nerve.

It has been reported that myelin debris is cleared by microglia and/or monocyte-derived macrophages in pathological conditions (Stoll et al. 1989; Stoll and Jander 1999; Weinger et al. 2011; Lloyd et al. 2017; Hammel et al. 2022). Recent studies have demonstrated that astrocytes (Wang et al. 2020; Hammel et al. 2022; Li et al. 2023) and microvascular endothelial cells (Zhou et al. 2019; Hammel et al. 2022; Yao et al. 2023) also clear myelin debris. These findings suggest that not only microglia and/or monocyte-derived macrophages but also astrocytes and microvascular endothelial cells phagocytose myelin debris-like MBPh-immunoreactive particles distributed in the m1ab region of the normal rat optic nerve. Further studies are required to determine whether this hypothesis is accurate or not.

In addition to the phagocytosis of myelin debris by microglia and/or by other phagocytes in pathological conditions, it has been demonstrated that myelin debris triggers an inflammatory response. For example, the direct application of myelin debris into the corpus callosum or the cerebral cortex induces profound and comparable inflammation in these structures of the mouse (Clarner et al. 2012; Grajchen et al. 2020; Hammel et al. 2022). It is conceivable that myelin debris-like MBPh-immunoreactive particles may also induce a slight inflammatory response in the m1ab region of the normal rat optic nerve. Further study is required to ascertain the accuracy of this hypothesis.

### 4.7 The physiologically stressed conditions in the m1ab region

The myelinated region of the normal rat optic nerve was previously recognized as homogeneously organized. Kawano (2015) demonstrated that the chemoarchitecture of this region was heterogeneously organized. In the present study, we confirmed these findings. We proposed that the heterogeneous distribution of the two glial cell marker proteins (GFAP, and GS) and of MBPh-immunoreactive particles in the myelinated region was attributable to the physiologically stressed conditions observed in the m1ab region of the normal rat optic nerve.

Although the m1ab region is part of the normal rat optic nerve, it is unclear why this region is under physiologically stressed conditions. One possible reason is that the m1ab region is located adjacent to the u (unmyelinated) region. The u region is one of the strongest regions of GFAP immunoreactivity (Kawano 2015). Consequently, the u region demonstrates a necessary requirement of gliosis: higher production of GFAP. Additionally, gliosis in the u region may be more severe due to the strongest GFAP immunoreactivity observed here. During reactive astrogliosis, cytokines, growth factors, and extracellular matrix proteins are released. These bioactive chemicals may be involved in immune response, neuroprotection, or potential further damage (Messam et al. 2002). It is possible that these bioactive chemicals are released in the u region and that they spread into the neighboring m1ab region. Consequently, the m1ab region may be slightly impaired not only by the bioactive chemicals released locally as a result of reactive astrogliosis (Section 4.4) but also by those originating from the adjacent u region.

This interpretation is supported by the fact that the capillary bed (Morrison et al. 1999; Sugiyama et al. 1999) is located in the proximal (posterior) half of the u region (Kawano 2015). The present study indicates the presence of capillary beds as defects in GFAP (Figure 1B) and GS immunoreactivity (Figure 2B) within this region. Since the capillary bed allows more nutrients and oxygen to enrich the tissue, thereby providing a healthier environment (Erickson and Liu-Ambrose 2016), it is assumed that the capillary bed is required in this region to prevent tissue impairment caused by reduced microvascular blood flow and the release of bioactive chemicals.

## 5 Conclusion

In summary, we investigated the distribution patterns of the glial cell marker proteins in the myelinated region of the normal rat optic nerve. GFAP and GS were distributed abundantly in the m1ab region, moderately in the m1cd region, and sparsely in both the m2ab and m2cd regions. Additionally, bBM-labeled cell nuclei were present plentifully in the m1ab region, and moderately in the m1cd, m2ab, and m2cd regions. Interestingly, MBPh-immunoreactive particles were distributed abundantly in the m1ab region but sparsely in the m1cd, m2ab, and m2cd regions.

Since reactive astrogliosis is characterized by a fibrous proliferation of glial cells in injured areas of the CNS (McAteer and Choudhury 2009; Sofroniew 2009), the abundant distribution of GFAP in the m1ab region suggests that this area might be under physiologically stressed conditions. Similarly, since GS in oligodendrocytes is increased in chronic pathological conditions in both mice and humans (Ben Haim et al. 2021), the abundant distribution of GS in the m1ab region also suggests that this area might be under physiologically stressed conditions. Additionally, since MBPh-immunoreactive particles are considered to be myelin debris-like structures (Kawano 2026), the abundant distribution of these particles in the m1ab region also suggests that this area might be under physiologically stressed conditions. Consequently, the concentration of GFAP, GS, cell nuclei, and of MBPh-immunoreactive particles in the m1ab region could represent significant functional adjustments that reflect the physiologically stressed state of the structural environment within this area of the normal rat optic nerve.

Oligodendrocyte myelination is regulated by axo-myelinic neurotransmission (Duncan et al. 2021). Glutamate functions as a key signaling molecule in relation to axo-myelinic neurotransmission in the normal rat optic nerve (Barron and Kim 2019). GS catalyzes the conversion of glutamate into glutamine, and a higher expression of GS in the m1ab region indicates a greater amount of vesicular glutamate release in this area (Kukley et al. 2007; Ziskin et al. 2007). Consequently, further studies on glutamate homeostasis in the optic nerve are necessary, particularly in the m1ab region, which is currently in a physiologically stressed state. These studies will enhance our fundamental understanding of the optic nerve and may provide new insights into various optic nerve diseases.

## Author Contributions

J.K.: Concept, Funding, Supervision, Investigation, Data analysis, Writing original draft, Writing – review and editing.

## Acknowledgments

The author thanks Professor Masahisa Horiuchi (Department of Hygiene and Health Promotion Medicine, Kagoshima University Graduate School of Medical and Dental Sciences) for his expert advice on statistical analyses. This work was supported by an annual fund from Kagoshima University.

## Data Availability Statement

The data that support the findings of this study are available from the corresponding author upon reasonable request.

## Patient Consent Statement

Not applicable.

## Permission to Reproduce Material from Other Sources

Not applicable.

## Clinical Trial Registration

Not applicable.

## Footnotes

1 In the companion pater (Kawano 2026), we used two types of anti-myelin basic protein (MBP) antibodies which detected different positions of the MBP protein in order to analyze distribution of the MBP protein in the normal rat optic nerve. One was an antibody that reacts with Ala-Ser-Asp-Tyr-Lys-Ser (ASDYKS) in position 131-136 of the classic human myelin basic protein (MBPh), while another was an antibody that reacts with Asp-Glu-Asn-Pro-Val-Val (DENPVV) in position 82-87 of the full length protein of cow myelin basic protein (MBPc). Here, we describe not MBP but MBPh to clarify which antibody was used to visualize the MBP protein.

2 To distinguish the m1 and m2 regions in the present study from those in the previous study (Kawano 2015), each subregion in the previous study is labeled with “*”: m1* and m2*.

3 In the companion paper (Kawano 2026), we used two types of anti-myelin basic protein (MBP) antibodies that recognize different positions of the MBP protein to analyze distribution of the MBP protein in the normal rat optic nerve. One was an antibody that reacts with Ala-Ser-Asp-Tyr-Lys-Ser (ASDYKS) in position 131-136 of the classic human myelin basic protein (MBPh). Another was an antibody that reacts with Asp-Glu-Asn-Pro-Val-Val (DENPVV) in position 82-87 of the full length protein of the cow myelin basic protein (MBPc). Here we clarify that we refer to MBPh rather than MBP itself to specify which antibody was used to visualize the MBP protein.

4 In our quantitative analyses, we used these modified images placed on a completely black background.

5 See footnote 3.

6 See footnote 3.

7 See footnote 3.

